# Phytoplankton performance in the lab predicts occurrence in the field across a global temperature gradient

**DOI:** 10.64898/2026.02.18.706362

**Authors:** Ting Lv, Fabio Benedetti, Dominic Eriksson, Meike Vogt, Mridul K. Thomas

## Abstract

Biologists aim to predict where species will survive and thrive as the planet warms. To do so, we often rely on data-hungry species distribution models (SDMs) that use associations between species occurrences and environmental predictors to capture the realised niche. An alternative basis for predictions is to experimentally quantify the effect of environmental drivers on performance, which captures the fundamental niche. We presently do not know which of these approaches represents a better path towards accurate forecasts. SDMs may depend too strongly on present-day environmental covariation, which will change in the future. In contrast, a major shortcoming of experiments is that they ignore most environmental drivers to focus on one or two. Quantifying how well fundamental and realised niches agree today would help establish how useful both SDMs and experiments are likely to be. We therefore compared both niches in 39 relatively common marine phytoplankton species. The temperature-dependence of population growth rate was characterised with a thermal performance curve model applied to lab experimental data, and the temperature-dependence of species occurrence probability estimated with SDMs applied to a global compilation of marine presence records. We found a fairly strong, near 1:1 relationship between measures of thermal niche centre: the *median growth temperature* in the lab and the *median occurrence temperature* in the field (R^2^ = 0.49). We also found a modest positive relationship between measures of thermal niche width, the *growth niche width* and the *occurrence niche width* (R^2^ = 0.24). This agreement should increase our confidence in environmental preferences inferred with SDMs. It also suggests that simple experiments can reliably constrain species ranges and help forecast range shifts. This has important implications for forecasting community composition and ecosystem processes, as we ought to be able to predict range shifts in biogeochemically-important taxa such as diatoms and nitrogen-fixing cyanobacteria.

## Introduction

Species distribution models (SDMs) capture statistical relationships between occurrence (or abundance) patterns and environmental predictors (Beaugrand et al., 2009; Elith and Leathwick, 2009). If one can forecast future environmental conditions, these relationships can be used to forecast future patterns of occurrence or abundance as well. However, extrapolation with flexible models is fraught with difficulties and we need to extrapolate outside the range of present-day conditions and predict species performance in future ‘no-analogue environments’ (Williams et al., 2007; Rivera et al., 2023). This extrapolation problem is grounded in the difference between the fundamental and the realised niche (Hutchinson, 1957; Pearman et al., 2008; Kearney and Porter, 2009; Matthiopoulos, 2022). The fundamental niche describes the range of environmental conditions that a species would survive in the absence of biotic interactions. The realised niche describes this range of conditions after its modification by biotic interactions (Hutchinson, 1957; Soberón, 2007). SDMs characterise the realized niche (often framed as ‘habitat suitability’) but this remains a challenge, particularly in the oceans. Despite substantial progress in SDM methods and data synthesis in the past decade (Buitenhuis et al., 2013; de Vargas et al., 2015; Sunagawa et al., 2020; Paoli et al., 2022; Valle et al., 2023; Vogt et al., 2023; Lombard et al., 2024; Balembois et al., 2025; Schickele et al., 2025), occurrence and abundance data remain scarce and highly biased. Data on environmental drivers is also often unavailable at the desired spatio-temporal scale. Modelled realised niches for important taxa such as plankton are therefore poorly constrained by observations (Dutkiewicz et al., 2020) and are almost never tested with experiments, limiting the degree to which we can trust them. Although developing forecasts based on fundamental niches characterised through controlled experiments seems like an appealing alternative, this has its own drawbacks. Experiments typically ignore intraspecific variation and measure just one or occasionally two dimensions of the niche, ignoring other drivers and biotic interactions, which are expected to interact in complex ways to shape species distributions.

Establishing that fundamental niches measured in lab experiments quantitatively predict realised niches in the field today (Bujan et al., 2022; Laeseke et al., 2024) would therefore serve two goals. It would establish the usefulness of measuring fundamental niches in lab experiments for predictive ecology. It would also bolster confidence in SDM-inferred species preferences, providing us with a concrete basis for forecasting range shifts and ecosystem changes. Despite intense interest in predicting these temperature-driven changes (Sunday et al., 2012; Benedetti et al., 2021a), this comparison has rarely been attempted, due to the challenge in both collecting occurrence data across broad spatial scales and in measuring the same species’ ecophysiological parameters in lab experiments (but see Guo et al., 2020; Camacho et al., 2024). We examine the match between fundamental and realised niches here across a wide range of species in marine phytoplankton, abundant and globally-distributed microbes that drive biogeochemical cycles and aquatic ecosystems (Falkowski et al., 1998; Field et al., 1998).

We focus here on a single dimension of the ecological niche, temperature. Temperature is probably the strongest abiotic driver of performance in most taxa (Brown et al., 2004; Beaugrand et al., 2008, 2009, 2013; Tittensor et al., 2010; Righetti et al., 2019; Benedetti et al., 2021a; Dal Bello and Abreu, 2024). We understand temperature–performance relationships well, both mathematically and empirically (e.g., Eppley, 1972; Ratkowsky et al., 1983; Huey and Hertz, 1984; Brown et al., 2004; Angilletta, 2009; Kingsolver, 2009; Huey et al., 2012; Thomas et al., 2012; Padfield et al., 2021). Growth curves (sometimes called thermal performance curves or TPCs; see Glossary in Table 1) characterise how performance changes across a temperature gradient. For microbes, we can parameterise established TPC functions that capture these left-skewed, unimodal relationships using data from simple lab growth assays and thereby characterise their fundamental thermal niches (Ratkowsky et al., 1983; Norberg, 2004).

**Table 1.** Glossary.

| Model-derived quantity | Data source used | Estimation / Calculation | Brief explanation (details in Methods) | Related terms |
| --- | --- | --- | --- | --- |
| Occurrence probability curve | Field presence-pseudoabsence and environmental predictors | Species Distribution Model (SDM) | A curve characterising how a species' probability of occurrence (i.e. presence) in the ocean changes as a function of temperature. It is the partial effect of temperature on occurrence probability, estimated from SDMs. Importantly, these probabilities are *uncalibrated*, meaning the *level* (height) of any curve is inaccurate. We focus on features of the curves' *shape* that should be robust. | Synonymous term in SDM literature: Response curve |
| Growth curve | Lab experiments measuring species' population growth rates at different temperatures | Thermal Performance Curve (TPC) model (Thomas et al., 2012) | A left-skewed unimodal curve characterising how a species' population growth rate changes as a function of temperature. This is characterised with a TPC model fitted to data from lab experiments measuring population growth rates at different temperatures. | Synonymous terms in thermal biology literature:<br>(i) Thermal performance curve (TPC)<br>(ii) Temperature response curve |
| Median occurrence temperature | Field presence-pseudoabsence and environmental predictors | Calculated from species' occurrence probability curve | The temperature at which the cumulative area under the occurrence probability curve reaches 50% of the total, in the temperature range [-1.8, 30]. In other words, 50% of the area under the curve occurs to either side of this temperature. |  |
| Median growth temperature | Lab experiments measuring species' population growth rates at different temperatures | Calculated from species' growth curve | The temperature at which the cumulative area under the growth curve reaches 50% of the total, in the temperature range [-1.8, 30] and the growth rate range of [0, ∞] i.e. we ignored negative growth rates. | Related term: Optimal temperature for growth |
| Occurrence niche width | Field presence-pseudoabsence and environmental predictors | Calculated from species' occurrence probability curve | The narrowest range of temperatures that contain 80% of the area under the occurrence probability curve, in the temperature range [-1.8, 30]. |  |
| Growth niche width | Lab experiments measuring species' population growth rates at different temperatures | Calculated from species' growth curve | The narrowest range of temperatures that contain 80% of the area under the growth curve, in the temperature range [-1.8, 30] and the growth rate range of [0, ∞] i.e. we ignored negative growth rates. | Related terms:<br>(i) Temperature niche width<br>(ii) Thermal niche breadth |

Temperature is especially important for phytoplankton. Both biogeochemical modelling and SDM studies consistently find it to be a major determinant of their spatial distribution (Dutkiewicz et al., 2013; Righetti et al., 2019; Benedetti et al., 2021a). It explains distributions of key taxa such as the nitrogen-fixer *Trichodesmium* (Karl et al., 2002; Monteiro et al., 2011; Litchman et al., 2012), as well as seasonal succession patterns at local scales (Karentz and Smayda, 1984). Growth curve parameters have been estimated for hundreds of species belonging to all major marine phytoplankton functional groups (Thomas et al., 2012, 2016), including coccolithophores (e.g., Buitenhuis et al., 2008; Jensen et al., 2017) and diatoms (e.g., Pinkernell and Beszteri, 2014; Jensen et al., 2017), enabling us to quantify fundamental thermal niches. Phytoplankton optimal temperatures for growth and other parameters are strongly related to mean sea surface temperatures at their isolation locations (Thomas et al., 2012, 2016), providing a strong basis for assuming that growth curves should predict realised niches as well.

However, these growth curves have not been quantitatively compared against field-based occurrence patterns across broad spatial scales, largely due to the absence of sufficient occurrence data. This has recently been addressed by the publication of global, cross-methods syntheses of phytoplankton species occurrences (e.g., Phytobase, MAREDAT, GBIF, and OBIS; Righetti et al., 2020; Benedetti et al., 2021b; Vogt et al., 2023). These occurrence records enable us to finally estimate the realised thermal niches of phytoplankton (Brun et al., 2015; Righetti et al., 2019; Benedetti et al., 2021a, 2023a). We describe these realised thermal niches here with SDM-derived ‘occurrence probability curves’ that characterise the nonlinear association between temperature and occurrence probability across the oceans (see Glossary in Table 1; Benedetti et al., 2021a; Eriksson et al., 2024). These probabilities are *uncalibrated* because we cannot entirely account for the underlying non-random sampling in the models. However, we believe that the features we focus on are robust to this (see Methods for details).

We investigated the relationship between species’ growth curves and their corresponding occurrence probability curves (Figs. 1, S1, S2). We compared proxies for optimum temperature and thermal niche width from both sets of curves. Specifically, we quantify how strongly (i) the *median growth temperature* in the lab is positively related to the *median occurrence temperature* in the field (Table 1), and (ii) the *growth niche width* in the lab is positively related to the *occurrence niche width* in the field (Table 1). We expect these two relationships to be broadly linear, with the possibility of nonlinearities near the extremes because of boundary conditions. This is because phytoplankton are short-lived organisms that are sensitive to environmental conditions, can rapidly grow when conditions are favourable (Furnas, 1990; Behrenfeld and Boss, 2018), and are not strongly affected by dispersal limitation (Jönsson and Watson, 2016; Villarino et al., 2018). Their populations are therefore likely to be approximately at equilibrium on the spatiotemporal scales considered in global SDMs. By integrating laboratory-derived ecophysiological data with field-based occurrence data, this work aims to bridge the gap between fundamental and realised niches, thereby advancing our understanding of how temperature shapes phytoplankton communities and informing models predicting how these communities will shift under future warming.

**Fig. 1.**
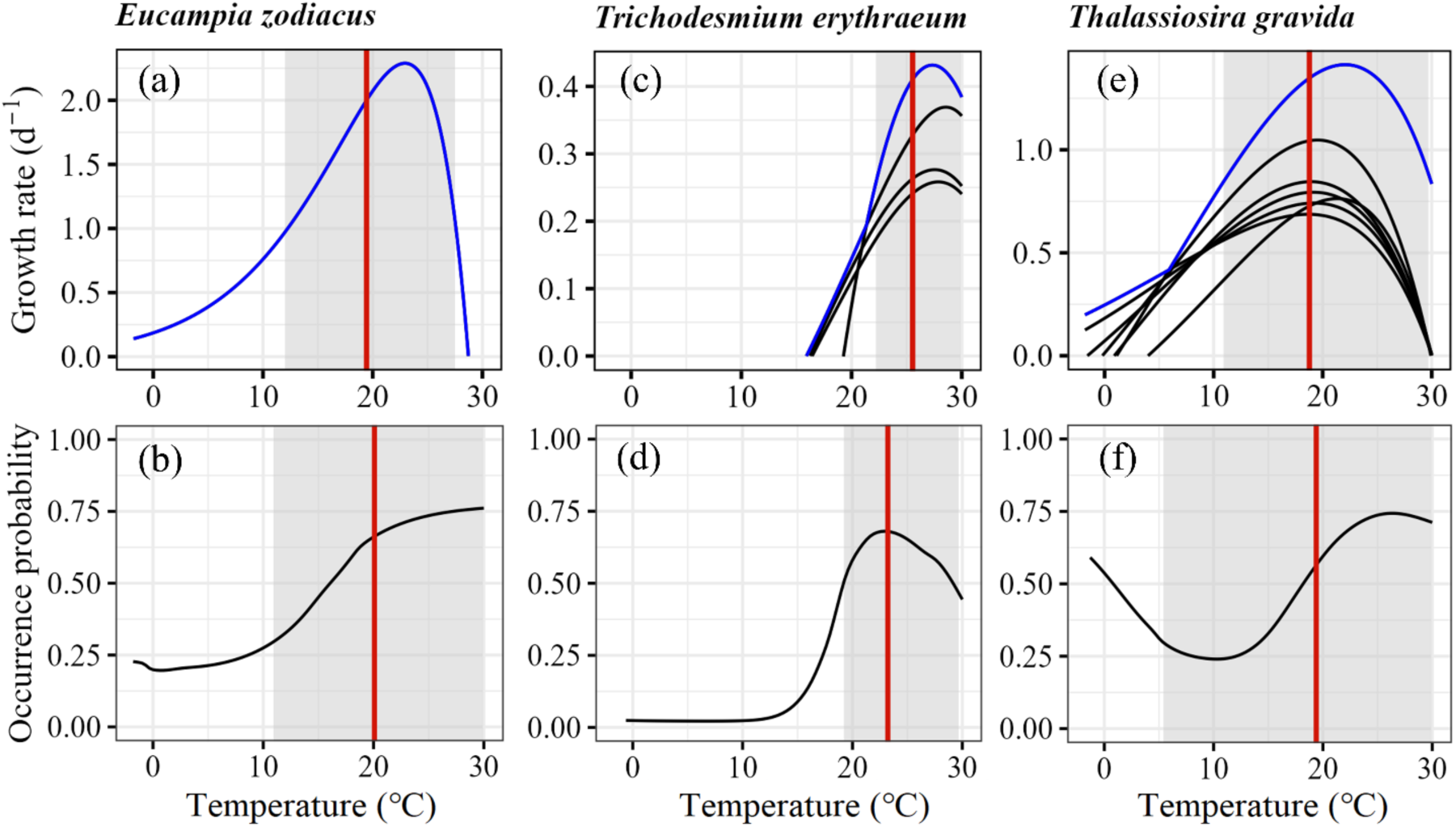
The temperature-dependence of population growth rate estimated from lab experiments (top row) and of uncalibrated occurrence probability estimated from SDMs fitted to presence/pseudoabsence data in the ocean (bottom row), for three marine phytoplankton species. These species are examples of the three shapes of occurrence probability curves that we manually classified curves into - monotonic, unimodal, and bimodal. We consider the few bimodal curves to be biologically unrealistic and exclude these species from our analyses. The vertical red lines show the median growth temperatures and median occurrence temperatures. The grey shaded regions indicated the growth niche widths and occurrence niche widths. For *Trichodesmium erythraeum* and *Thalassiosira gravida*, different growth curves (in black) correspond to different published experimental datasets, largely representing different strains. We overlay the *envelope* of the individual curves in blue and use this to calculate the indicated median growth temperature and growth niche width. Curves for all species in this study are shown in Figs. S1 and S2.

**Fig. 2.**
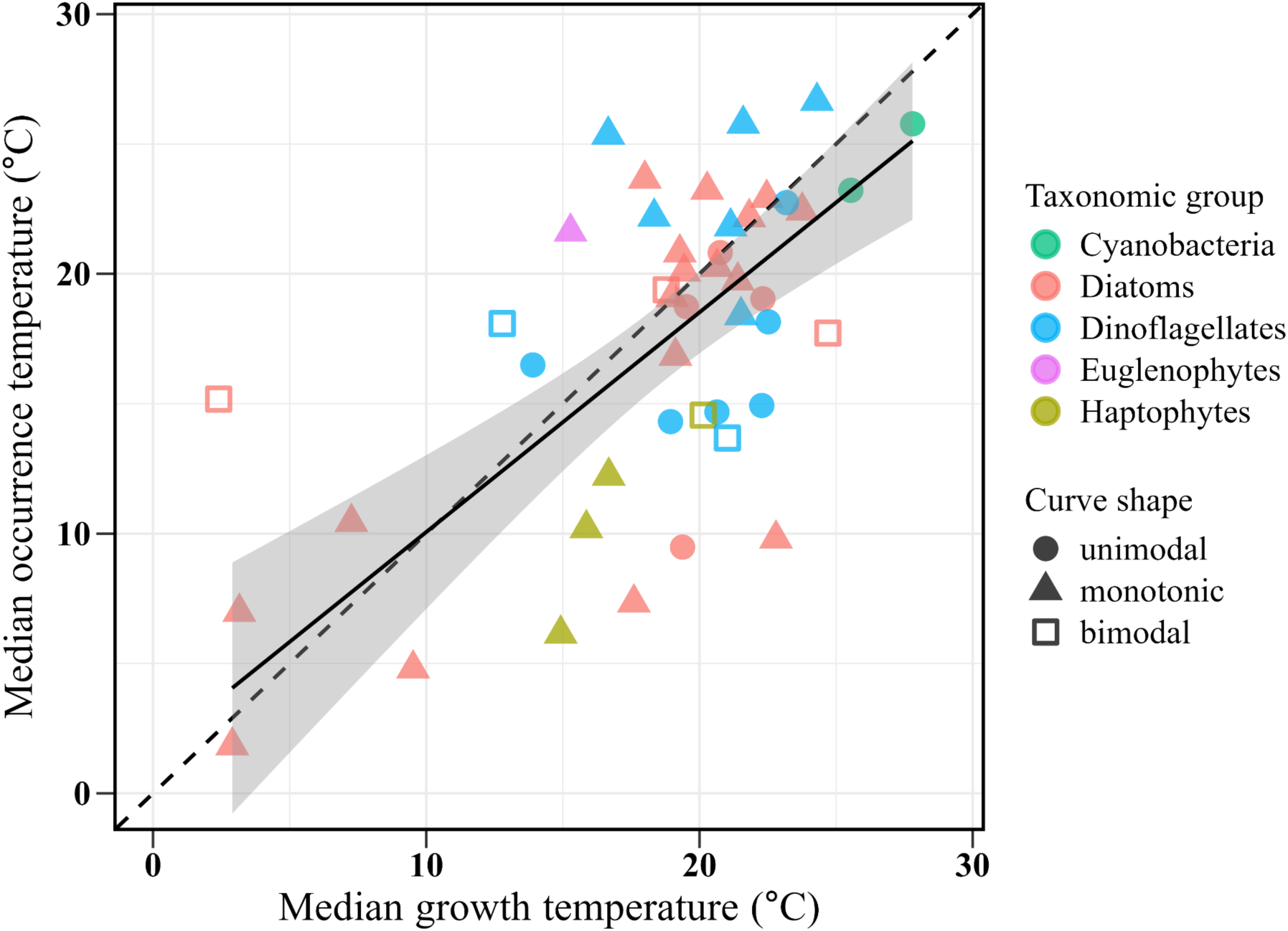
Median growth temperature in the lab predicts median occurrence temperature in the field reasonably well across 39 marine phytoplankton species (slope = 0.85, intercept = 1.60, R^2^ = 0.49, *p* < 0.001, Table S2). The mean response (solid regression line) is close to the 1:1 line (dashed line). Median occurrence temperature estimates are likely most reliable for unimodal curves, followed by monotonic and finally bimodal curves. The six bimodal curves (hollow squares) were therefore excluded from the regression. Where lab growth curves exist for multiple strains of the same species, points are calculated based on the envelope of all strains’ growth curves.

## Methods

### Overview

We used compilations of phytoplankton thermal performance curve parameters (Thomas et al., 2012, 2016) and observations of field occurrences (Righetti et al., 2020; Eriksson et al., 2024) to estimate how strongly lab growth curves predict thermal preferences inferred from species distribution models. We focus on 45 species belonging to 5 plankton functional groups for which (i) there was a published lab growth curve, and (ii) we had at minimum 75 presence records in the field data (Benedetti et al., 2021a). The species, many of which are relatively common and broadly-distributed, are listed in Table 2. The occurrences for these 45 species comprise 15.7% of all diatom records, 22.6% of dinoflagellate records, 46.2% of haptophyte records, 96.5% of euglenophyte records, and 9.5% of cyanobacteria records (we did not analyse *Prochlorococcus* and *Synechococcus* as they are not identified to a species level). Note that these percentage calculations omit unidentified individuals. All lab growth curves and field occurrence probability curves used in this paper are shown in the supplementary information (Figs. S1, S2). We then performed linear regressions of median growth temperature against median occurrence temperature, and growth niche width against occurrence niche width.

**Table 2.** Summary of the 45 focal species with data available from both laboratory experiments and field observations. Three distinct occurrence curve shapes were identified: unimodal, monotonic, and bimodal (note that we omitted bimodal curves from all models).

| Taxonomic group | Species | Shape of occurrence probability curve | Median growth temperature | Median occurrence temperature | Growth niche width | Occurrence niche width |
| --- | --- | --- | --- | --- | --- | --- |
| Diatoms | *Asterionellopsis glacialis* | unimodal | 19.5 | 18.7 | 18.7 | 19.5 |
|  | *Chaetoceros affinis* | monotonic | 21.8 | 22.2 | 12.1 | 13.4 |
|  | *Chaetoceros didymus* | monotonic | 21.4 | 19.7 | 12.8 | 19.8 |
|  | *Chaetoceros lorenzianus* | monotonic | 22.5 | 22.9 | 13.8 | 12.1 |
|  | *Chaetoceros pseudocurvisetus* | monotonic | 23.8 | 22.4 | 9.9 | 12.8 |
|  | *Chaetoceros simplex* | monotonic | 22.8 | 9.8 | 11.8 | 22.9 |
|  | *Corethron pennatum* | bimodal | 2.4 | 15.2 | 6.1 | 25.5 |
|  | *Coscinodiscus concinnus* | monotonic | 17.6 | 7.3 | 13.9 | 19.7 |
|  | *Coscinodiscus granii* | monotonic | 18.0 | 23.7 | 17.1 | 12.3 |
|  | *Coscinodiscus wailesii* | monotonic | 19.3 | 20.8 | 19.0 | 18.3 |
|  | *Dactyliosolen fragilissimus* | monotonic | 19.0 | 19.1 | 16.2 | 20.1 |
|  | *Detonula confervacea* | monotonic | 7.3 | 10.4 | 12.5 | 22.9 |
|  | *Ditylum brightwellii* | unimodal | 20.8 | 20.8 | 17.0 | 15.9 |
|  | *Eucampia zodiacus* | monotonic | 19.4 | 20.1 | 15.5 | 19.0 |
|  | *Fragilariopsis kerguelensis* | monotonic | 2.9 | 1.8 | 6.9 | 10.7 |
|  | *Helicotheca tamesis* | unimodal | 22.3 | 19.0 | 12.6 | 21.7 |
|  | *Leptocylindrus danicus* | bimodal | 24.7 | 17.7 | 9.4 | 23.9 |
|  | *Proboscia inermis* | monotonic | 3.2 | 7.0 | 6.3 | 16.9 |
|  | *Rhizosolenia setigera* | monotonic | 20.7 | 20.3 | 17.7 | 19.7 |
|  | *Skeletonema costatum* | monotonic | 19.1 | 16.8 | 18.6 | 23.3 |
|  | *Stephanopyxis palmeriana* | monotonic | 20.3 | 23.2 | 14.0 | 11.1 |
|  | *Thalassionema nitzschioides* | unimodal | 19.4 | 9.5 | 20.0 | 18.8 |
|  | *Thalassiosira gravida* | bimodal | 18.8 | 19.4 | 18.8 | 24.5 |
|  | *Thalassiosira nordenskiöldii* | monotonic | 9.5 | 4.8 | 14.4 | 12.6 |
| Haptophytes<br>(coccolithophores<br>+ *Phaeocystis*) | *Calcidiscus leptoporus* | monotonic | 16.7 | 12.2 | 11.6 | 23.8 |
|  | *Coccolithus pelagicus* | monotonic | 15.9 | 10.2 | 14.8 | 21.8 |
|  | *Emiliania huxleyi* | bimodal | 20.1 | 14.6 | 18.1 | 24.8 |
|  | *Phaeocystis pouchetii* | monotonic | 15.0 | 6.1 | 11.2 | 13.2 |
| Cyanobacteria | *Crocospaera watsonii* | unimodal | 27.8 | 25.8 | 4.1 | 7.2 |
|  | *Trichodesmium erythraeum* | unimodal | 25.5 | 23.2 | 7.7 | 10.5 |
| Dinoflagellates | *Akashiwo sanguinea* | unimodal | 20.6 | 14.7 | 16.5 | 23.9 |
|  | *Dinophysis caudata* | unimodal | 23.2 | 22.7 | 13.7 | 11.7 |
|  | *Leonella granifera* | monotonic | 24.3 | 26.7 | 10.1 | 7.0 |
|  | *Prorocentrum balticum* | bimodal | 21.0 | 13.7 | 13.2 | 26.7 |
|  | *Prorocentrum cordatum* | monotonic | 21.5 | 18.4 | 13.6 | 21.7 |
|  | *Prorocentrum gracile* | monotonic | 16.7 | 25.4 | 10.3 | 9.1 |
|  | *Prorocentrum lima* | monotonic | 18.3 | 22.2 | 18.5 | 16.5 |
|  | *Prorocentrum micans* | monotonic | 21.1 | 21.8 | 13.6 | 16.4 |
|  | *Scrippsiella acuminata* | bimodal | 12.8 | 18.1 | 17.6 | 26.4 |
|  | *Thoracosphaera heimii* | monotonic | 21.6 | 25.8 | 14.1 | 12.8 |
|  | *Tripos furca* | unimodal | 22.5 | 18.2 | 12.4 | 19.1 |
|  | *Tripos fusus* | unimodal | 22.3 | 14.9 | 14.2 | 22.6 |
|  | *Tripos lineatus* | unimodal | 18.9 | 14.3 | 15.0 | 21.8 |
| Euglenophytes | *Eutreptiella gymnastica* | monotonic | 15.3 | 21.6 | 17.5 | 18.0 |

### Estimating parameters of growth curves based on lab experiments

#### Data sources

Thomas et al. (2012) and (2016) compiled published lab measurements of phytoplankton population growth rates across a range of temperatures. After quality control, the dataset included measurements from 249 marine strains belonging to 115 species. We refer readers to these studies for details. We additionally extracted the data for two further species (*Dinophysis caudata*, *Prorocentrum balticum*) from more recently published studies (Basti et al., 2015; Funaki et al., 2024).

#### Thermal performance curve (TPC) model fitting

For each strain in the dataset, Thomas et al. (2012) and (2016) used the compiled data to estimate the parameters of a TPC function adapted from Norberg (2004):

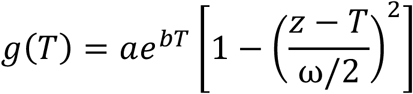

Here, the specific growth rate *g* depends on temperature *T*. Parameters z and ω determine the location and scale of the growth curve, respectively. *a* and *b* jointly determine the curve’s height and skewness. Further details may be found in Thomas et al. (2016).

We obtained curve parameters for 17 additional growth curves belonging to six species (*Emiliania huxleyi, Chaetoceros simplex*, *Coscinodiscus concinnus*, *Coscinodiscus granii*, *Prorocentrum lima* and *Thoracosphaera heimii*) from an updated version of this growth curve database (Kremer, personal communication). *E. huxleyi* was already represented in our dataset but this added 12 new strains. For the two species from more recent studies whose data we extracted for this paper (*Dinophysis caudata*, *Prorocentrum balticum*), we estimated their curve parameters ourselves following the same workflow used in Thomas et al. (2012) and Thomas et al. (2016).

#### Parameter estimation for this study

Using these TPC parameters, we recreated the growth curves for every strain of the focal 45 species (Figs. 1, S1). We estimated a proxy for the optimum temperature for growth that we term the *median growth temperature*. We estimated this in the temperature range [−1.8, 30], which reflects the approximate range of ocean temperatures in the field dataset (Toseland et al., 2013; Righetti et al., 2019), and the growth rate range of [0, ∞] i.e. we ignored negative growth rates. We then calculated the cumulative area under each growth curve as a function of temperature by numerically integrating the growth rates with respect to temperature using a step size of 0.02 ℃ and spline interpolation (using base R functions *integrate* and *splinefun*). The cumulative area percentage was obtained by dividing the cumulative area at each temperature point by the total area under the curve. We defined the *median growth temperature* to be the temperature at which this cumulative area percentage reached 50%. This median growth temperature is quantitatively extremely similar to the optimum temperature for growth (Fig. S4). We nonetheless chose to use *median growth temperature* to be consistent with the methods we needed to use for the field occurrence probability curves.

Secondly, we estimated the narrowest range of temperatures that contain 80% of the area under the curve and termed this the (thermal) *growth niche width*. We note that this is not identical to the thermal niche breadth - a term occasionally used in the thermal biology literature - which quantifies the range of temperatures over which growth rate is at least 80% (or some other percentage) of the maximum growth rate. To quantify the growth niche width, we systematically evaluated all possible temperature intervals within each species’ growth curve, using a step size of 0.1 ℃. For each interval, we again calculated the area under the curve using numerical integration with spline interpolation. Among this set of intervals, we selected those where the area equalled 80% of the total area under the curve, and defined the narrowest interval in this set as the growth niche width.

For a subset of species, we have TPC parameters for multiple strains in the same species. However, field occurrence data only occurs at the species level (if that). We therefore aggregated lab estimates to the species level to make the two datasets comparable. We did this by estimating the median growth temperature and the growth niche width from the *envelope* of all strains’ growth curves (Figs. 1, S1). That is, we assume that the fastest growing strain of every species will dominate at any temperature. For each species, we first use each strain’s growth curve to predict the growth rate in the interval [−1.8, 30] at an interval of 0.02 ℃. We then calculate the maximum growth rate across strains at each temperature interval and fit a spline to this maximum growth rate. The median growth temperature and the growth niche width were calculated on this spline as previously described. As an additional analysis, we separately estimated these parameters for each strain of every species and took the mean within species to define a species’ parameters. This method produced similar results (Figs. S6, S9). We also used these strain-level estimates to evaluate the relationships between median growth temperature, growth niche width, and isolation location.

### Estimating parameters of occurrence probability curves based on field observations

We used the niche modelling approach described in Benedetti et al. (2021a) and Benedetti et al. (2023a) to estimate the occurrence probability curves of the 45 target species.

#### Data sources

We used phytoplankton species occurrence data compiled in Righetti et al. (2020) and Eriksson et al. (2024). Righetti et al. (2020) integrates data from several key sources: the Global Biodiversity Information Facility (GBIF; https://www.gbif.org), the Ocean Biogeographic Information System (OBIS; https://www.obis.org), Villar et al. (2015), and the MARine Ecosystem biomass DATa (MAREDAT) initiative (Buitenhuis et al., 2013). Species nomenclature was standardized and curated in accordance with the taxonomic framework provided by AlgaeBase (http://www.algaebase.org/). This dataset encompasses over one million individual records, representing approximately 1700 phytoplankton species, collected using a range of sampling methodologies. All records are situated within the monthly climatological mixed layer depth, with a sampling depth of 5.4 ± 6.9 m (mean ± SD), spanning the period from 1800 to 2015. We further used field records gathered by Eriksson et al. (2024) on marine surface planktonic nitrogen-fixing microorganisms to model two target nitrogen-fixers (*Crocosphaera watsonii* and *Trichodesmium erythraeum*) that could not be investigated with the data of Righetti et al. (2020). The nitrogen fixer database contained approximately 22,000 records gathered from GBIF (last access: 20 October 2021), OBIS (last access: 21 October 2021), Luo et al. (2012), Tang and Cassar (2019), Gradoville et al. (2020), Righetti et al. (2020), Pierella Karlusich et al. (2021), Detoni et al. (2022), Martinez-Gutierrez et al. (2023), and the Ocean Microbiomics Database (Paoli et al., 2022), using either microscopy-based or sequence-based (qPCR or metagenomics) sampling methodologies across all ocean basins, with a mean sampling depth of 36 ± 47 m (mean ± SD), spanning the period from 1880 to 2021.

#### Species Distribution Model (SDM) fitting

We modelled occurrence (i.e., presence) probability as a function of six environmental predictors identified as important for phytoplankton species distribution modelling by Benedetti et al. (2021a, 2023a): (i) monthly mean sea surface temperature (SST, °C), (ii) photosynthetically active radiation (PAR, μmol m^−2^ s^−1^), (iii) surface concentration of silicic acid Si(OH)_4_ (μM, on a log scale) (iv) the excess of Si(OH)_4_ relative to nitrate NO_3_^−^ (Si^∗^ = Si(OH)_4_ - NO_3_^−^, μM) (v) excess of nitrate to phosphate relative to the Redfield ratio (N^∗^ = NO_3_^−^ - 16∗PO_4_^3−^, μM), and (vi) surface phytoplankton chlorophyll-a concentration (mg m^−3^, on a log scale). SST and the four nutrient-related fields were sourced from the World Ocean Atlas 2013v2 (Boyer et al., 2013; https://www.nodc.noaa.gov/OC5/woa13/woa13data.html). Surface phytoplankton chlorophyll-a concentration - a proxy for primary productivity here - was sourced from the merged case 1 water product from the GlobColour project (Maritorena et al., 2010; https://hermes.acri.fr/). All environmental layers were formatted as 12 global monthly climatologies following the standard 1 × 1 global cell grid of the WOA. Previous studies identified these six predictors as useful in predicting phytoplankton species’ niches based on univariate importance rankings and permutation-based tests (Brun et al., 2015; Barton et al., 2016; Ajani et al., 2018; Righetti et al., 2019; Benedetti et al., 2023a). All modelled species had at least 75 occurrence records, allowing us to maintain an occurrence-to-predictor ratio exceeding 12:1 — above the 10:1 threshold recommended by Guisan et al. (2017). To avoid collinearity-related biases in SDM projections (Dormann et al., 2013), pairwise Spearman’s rank correlations (ρ) were examined within each species’ dataset to make sure that the present pairs of predictors all display a median |ρ| < 0.75. A more exhaustive description of the environmental predictors tested and their relative predictive power can be found in Benedetti et al. (2023a). We underline that SST was consistently found to be the top-ranked predictor for the groups represented by our 45 target species (Benedetti et al., 2023a; Eriksson et al., 2024).

To provide the SDMs with information about the environmental space unused by the target species, ‘background data’ (i.e., pseudo-absences; Barbet-Massin et al., 2012) was generated using the target-group method (Phillips et al., 2009). This method samples points from locations where species from the same plankton functional group were observed but the focal species was not, and treats these points as locations where the species was absent. This approach: (i) minimizes sampling bias, (ii) avoids misclassifying unsampled areas as unsuitable. For each species, pseudo-absences were drawn at a 10:1 ratio relative to presences and weighted inversely to presence frequency (Barbet-Massin et al., 2012). This approach, standard within the SDM field, necessarily means that the occurrence probability curves are not well-calibrated mathematical probabilities (which is why they are sometimes referred to as ‘habitat suitability indices’). This means that the quantitative values of probability indicated in Fig. 1 and S2 are not representative of their absolute occurrence probability, i.e. a value of 0.5 does not imply a 50% probability of finding the species there. Addressing this limitation would require random or stratified sampling at a scale that was not feasible in this study, along with either true absences or completely unbiased pseudoabsences sampled at the correct frequency. While this influences the overall elevation of the occurrence–probability curves, it is unlikely to substantially affect their shape or the estimates of median occurrence temperature. Estimates of occurrence niche width, however, may be subject to a modest bias.

We fit three types of SDMs of varying complexity to the data from each species: generalized linear models (GLMs), generalized additive models (GAMs), and artificial neural networks (ANNs). To mitigate risks of overfitting (Merow et al., 2014), we limited the number of environmental predictors in relation to the number of occurrence records (see below) and constrained the models to yield simple response functions. Each SDM was run ten times with a five-fold cross validation strategy and the resulting True Skill Statistic (TSS) and Area Under the Curve (AUC) were calculated to evaluate SDM skill. The species presence/pseudo-absence data were randomly split into ten training and testing sets (80–20%, respectively) and 30 models (three SDM types × ten cross-validation sets) were trained per target species. A comprehensive account of model parameterization, including an assessment of sensitivity to sampling biases and input data characteristics, can be found in Benedetti et al. (2021a, 2023a).

#### Parameter estimation for this study

We extracted the partial dependence of occurrence on temperature for every species from their 30 SDM ensemble members. This partial dependence represents the nonlinear association between climatological monthly mean SST and occurrence probability, estimated from the model ensemble when all other predictors are set at their mean global surface value in the dataset. Specifically, it represents the mean across the ensemble members. We term this the species’ occurrence probability curve. Out of the 45 species occurrence probability curves, 12 showed the expected unimodal shape, while 27 cases were monotonic and 6 were bimodal (*Emiliania huxleyi*, *Corethron pennatum*, *Leptocylindrus danicus, Prorocentrum balticum, Scrippsiella acuminata and Thalassiosira gravida*). *Akashiwo sanguinea*’s curve was ambiguous (Fig. S2); we chose to classify it as unimodal. We expect that the monotonic curves largely reflect data limitations in the tropics and poles, with most occurrences and pseudoabsences being located at temperate latitudes (Graham et al., 2008; Hofner et al., 2011).

Following the procedure we used with the lab curves, we then estimated a proxy for every species’ optimum temperature in the field that we term the *median occurrence temperature*. For each species, this is the temperature at which the cumulative area under the occurrence probability curve reached 50% of the total in the temperature range [−1.8, 30]. In other words, 50% of the area under the curve occurs to either side of this temperature (Figs. 1, S2). In contrast to the strong linear relationship between median growth temperature and the temperature of maximum growth rate (i.e. optimum temperature for growth), the median occurrence temperature exhibits a sigmoidal relationship with the temperature of maximum occurrence probability (Fig. S5). The bimodal distribution of the temperature of maximum occurrence probability, which results from the majority of the curves being monotonic, contains little useful ecological information and motivated us to develop the median occurrence temperature metric.

We also estimated the narrowest range of temperatures that contains 80% of the area under the curve, and term this the *occurrence niche width*. As with the *growth niche width*, we systematically evaluated all possible temperature intervals within each species’ occurrence probability curve, using a step size of 0.1℃. For each interval, we calculated the area under the curve using numerical integration with spline interpolation. Among these intervals, we identified those where the area equalled 80% of the total area under the curve and defined the one with the narrowest temperature range as the *occurrence niche width*. This area-based metric provides an integrative measure of performance or suitability across the thermal gradient, while reducing sensitivity to poorly constrained range margins and associated edge effects in SDM-derived response curves (Elith and Leathwick, 2009; Monahan, 2009; Smith et al., 2021). To evaluate the robustness, we performed a sensitivity analysis by testing alternative thresholds from 40% to 90% at 10% increments. The results were consistent across thresholds, with the variance explained being highest at the 70% and 80% thresholds (R² = 0.24; Fig. S3). The 80% threshold we used was chosen before this robustness analysis.

We include all parameter estimates from both growth and occurrence probability curves (which are shown in Figs. S1, S2) in the supplementary information.

#### Methodological note

We evaluate occurrence probability curves (and therefore growth curves) over the temperature interval [−1.8, 30], which is the realized global range of surface ocean monthly mean temperatures in today’s ocean. Altering this range will alter the estimated median occurrence temperatures even without a change in the physical or biological data. Due to the preponderance of monotonic curves, the conceptually preferable ‘temperature of peak occurrence probability’ (analogous to the optimum temperature for growth) exhibits an uninformative bimodal distribution concentrated at the temperature extremes (Fig. S5). This may change with better presence/pseudo-absence data from the tropics and poles but at present, we believe that median occurrence temperature is more informative. We note that a similar approach is applied to estimate niche centroids in the SDM literature (Monahan and Tingley, 2012; Smith et al., 2021; Zhu et al., 2024).

### Statistical analyses

We used linear regressions (based on ordinary least squares, OLS) to compare median growth and occurrence temperatures, and also growth and occurrence niche widths. Because the niche width model’s residuals deviated moderately from normality, we re-fit the regressions with robust regression techniques using Iteratively Reweighted Least Squares (IRLS), with both Huber and bisquare weights. As parameter estimates and uncertainties were quantitatively very similar, we present the result from the original OLS regressions. We also explored whether differences between median growth and occurrence temperatures are associated with latitude, growth niche width, or maximum growth rate. For latitude, we used linear mixed models with a random intercept term to account for non-independence due to our use of strain-level parameter estimates, as strains from the same species were isolated from different locations. We used parametric bootstrapping with 10,000 bootstraps to estimate *p*-values for this mixed model. For growth niche width and maximum growth rate, we used linear regressions with the species-level parameter estimates based on the envelope of all strains’ growth curves.

### Software tools used

All data analyses and numerical integrations were performed using R version 4.4.2 (R Core Team, 2024). We used the following packages: *biomod2* for fitting SDMs (Thuiller et al., 2023), *snakecase* for formatting species names (Grosser, 2023), *dplyr* for data manipulation (Wickham et al., 2023), *lme4* for mixed model parameter estimation (Bates et al., 2015), *pbkrtest* for parametric bootstrapping of mixed models (Halekoh and Højsgaard, 2014), *MASS* for robust regression (Venables and Ripley, 2002), *sjPlot* for generating tables summarising fitted models (Lüdecke, 2024), and *ggplot2* (Wickham, 2016) for plotting.

## Results

We compared the fundamental and realised thermal niches for 45 marine phytoplankton species comprising 24 diatoms, 4 haptophytes (comprising coccolithophores + *Phaeocystis*), 14 dinoflagellates, 2 cyanobacteria and 1 euglenophytes (Table 2, Figs. S1, S2). Of these, we excluded 6 species from our analyses because their occurrence probability curves were bimodal, which we believe reflects data or modelling limitations. Note that the species list does not constitute a random sample of marine phytoplankton, as not all phytoplankton can be easily grown in the lab, identification is less reliable for rarer and smaller species in field samples, and taxonomic uncertainties abound.

### Temperature-dependent growth and occurrence probability curves

The occurrence probability curves exhibited three distinct shapes: unimodal, monotonic, and bimodal. Just 12 of the 45 species’ occurrence probability curves showed the expected unimodal shape (Fig. 1d, Table 2), while 27 were monotonic (either increasing or decreasing, Fig. 1b, Table 2) and 6 were bimodal (*Emiliania huxleyi*, *Corethron pennatum*, *Leptocylindrus danicus, Prorocentrum balticum, Scrippsiella acuminata and Thalassiosira gravida*; Fig. 1f, Table 2). Though we expected to see unimodal responses, a monotonic response is not unreasonable over the range of ocean temperatures, as tropical or polar species are, by definition, most likely to be found at one extreme of the temperature range. The bimodal curves that we excluded from our analyses may have more complex underlying causes.

### Median growth temperature vs. median occurrence temperature

Across the 39 species (excluding the 6 species with bimodal occurrence probability curves), the median growth temperature estimated from lab experiments was reasonably strongly associated with median occurrence temperature in the field (Fig. 2, R^2^ = 0.49, *p* < 0.001, Table S2). The modelled mean response closely follows the 1:1 line (slope = 0.85, intercept = 1.60; concordance correlation coefficient = 0.61). There were however ten species whose median growth and occurrence temperatures differed by more than 5°C, two of which differed by more than 10°C (*Chaetoceros simplex* and *Coscinodiscus concinnus*). The quantity of the occurrence data and underlying quality of the species distribution models - whether measured by AUC or TSS - do not appear to explain variation around the regression line (Fig. S6). We see no clear evidence for differences in the regression relationship between groups. Regression results remained extremely similar when the species’ median growth temperatures were calculated as the mean of the estimates of the individual strains, instead of being calculated from the envelope (Fig. S7).

When analysed at the strain - instead of species - level, the difference between median growth temperature and median occurrence temperature displays a modest geographical signal. This difference decreases weakly with absolute latitude (marginal R^2^ = 0.18, Fig. 3, Table S3). In other words, tropical strains have estimated median growth temperatures that are higher than their median occurrence temperatures, while for temperate strains they are similar or slightly below. This may reflect temporal specialisation of different strains at temperate latitudes, where there is more thermal variation. It appears to be unrelated to other aspects of the growth curves; the difference exhibits no meaningful association with growth niche width (Fig. S8, *p* = 0.78) or maximum growth rate (Fig. S9, *p* = 0.45).

**Fig. 3.**
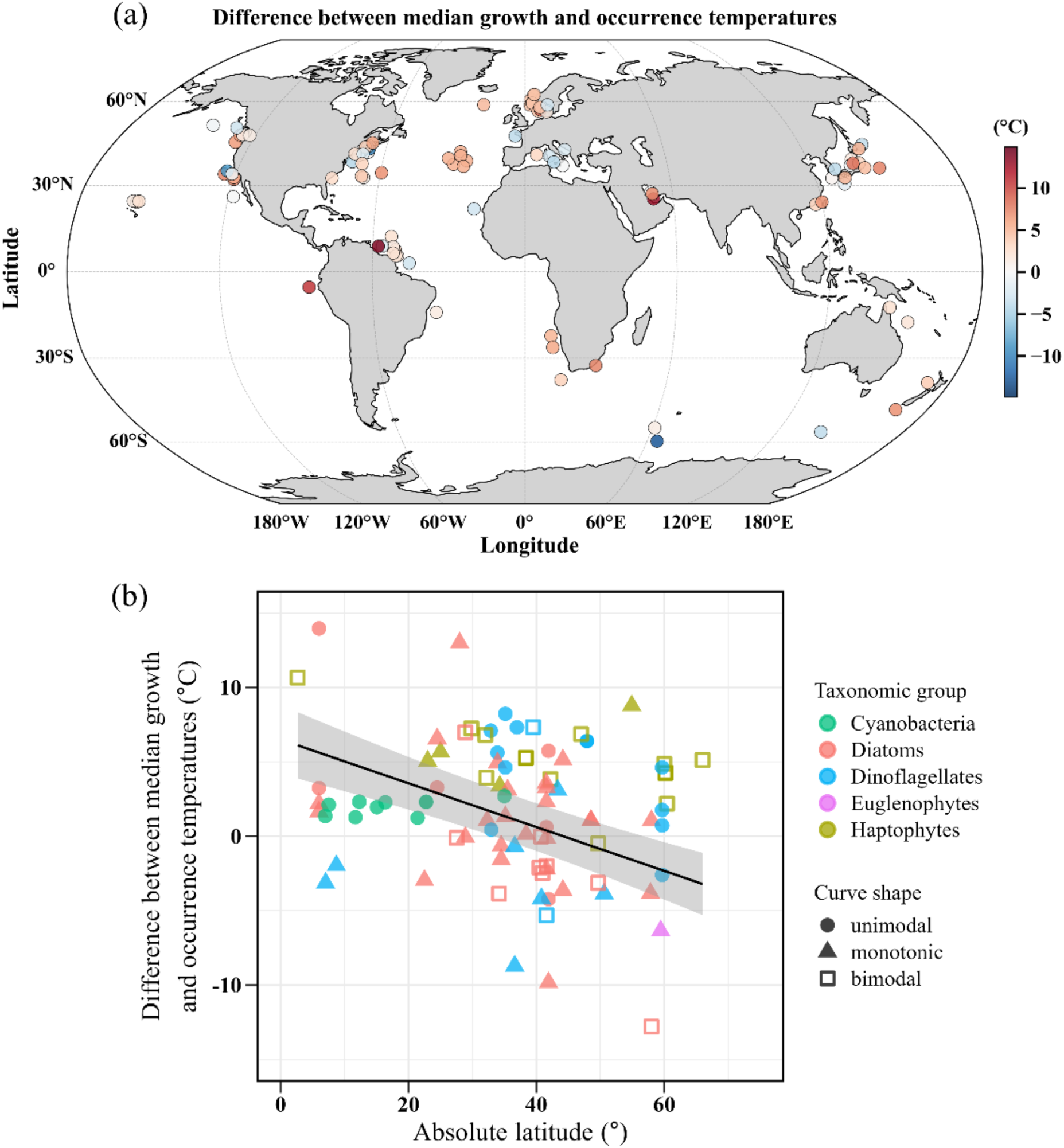
At the strain (instead of species) level, the difference between the median growth and occurrence temperatures changes with latitude. Positive values indicate higher median growth temperatures than median occurrence temperatures. Each point represents a phytoplankton strain with a measured lab growth curve at its isolation location. Unlike in Fig. 2, we estimated median growth temperatures separately for each of the 97 strains belonging to 41 species, because for some species, we have growth curves measured for multiple strains with different isolation locations. Note that there are four additional species and multiple strains in our growth curve dataset that are not represented here because we do not know their isolation locations. (a) Geographic variation in the difference between median growth and median occurrence temperatures. (b) The difference between median growth and median occurrence temperatures decreases weakly with absolute latitude (slope = −0.15, bootstrap *p* = 0.001, marginal R^2^ = 0.18, Table S3). The regression parameters were estimated using a mixed-effects model with a random intercept term to account for non-independence due to having multiple strains from the same species. Of the 97 points shown, 65 were used to fit the model; the 32 excluded points belonged to the three species with bimodal occurrence probability curves.

### Growth niche width vs. occurrence niche width

We estimated niche widths from growth and occurrence curves as the narrowest range of temperatures that contained 80% of the area under the curve in the interval [−1.8, 30]. Consistent with our expectations, these growth and occurrence niche widths were positively associated but the relationship was of modest strength (Fig. 4, slope = 0.62, R^2^ = 0.24, *p* = 0.002, Table S4). The residual distribution deviates moderately from a Gaussian distribution but re-analysis using robust regression leads to nearly identical coefficients and standard errors, so we present the original estimates here. Results are similar if species’ growth niche widths are calculated as the mean of the estimates of the individual strains, instead of being calculated from the envelope (Figs. 4, S10). Occurrence niche widths were generally higher, whether their occurrence probability curves were monotonic or unimodal. This indicates a higher probability of occurrence at high temperatures than is predicted based on the growth curves, reflecting the fact that occurrence curves were generally flatter than lab growth curves.

**Fig. 4.**
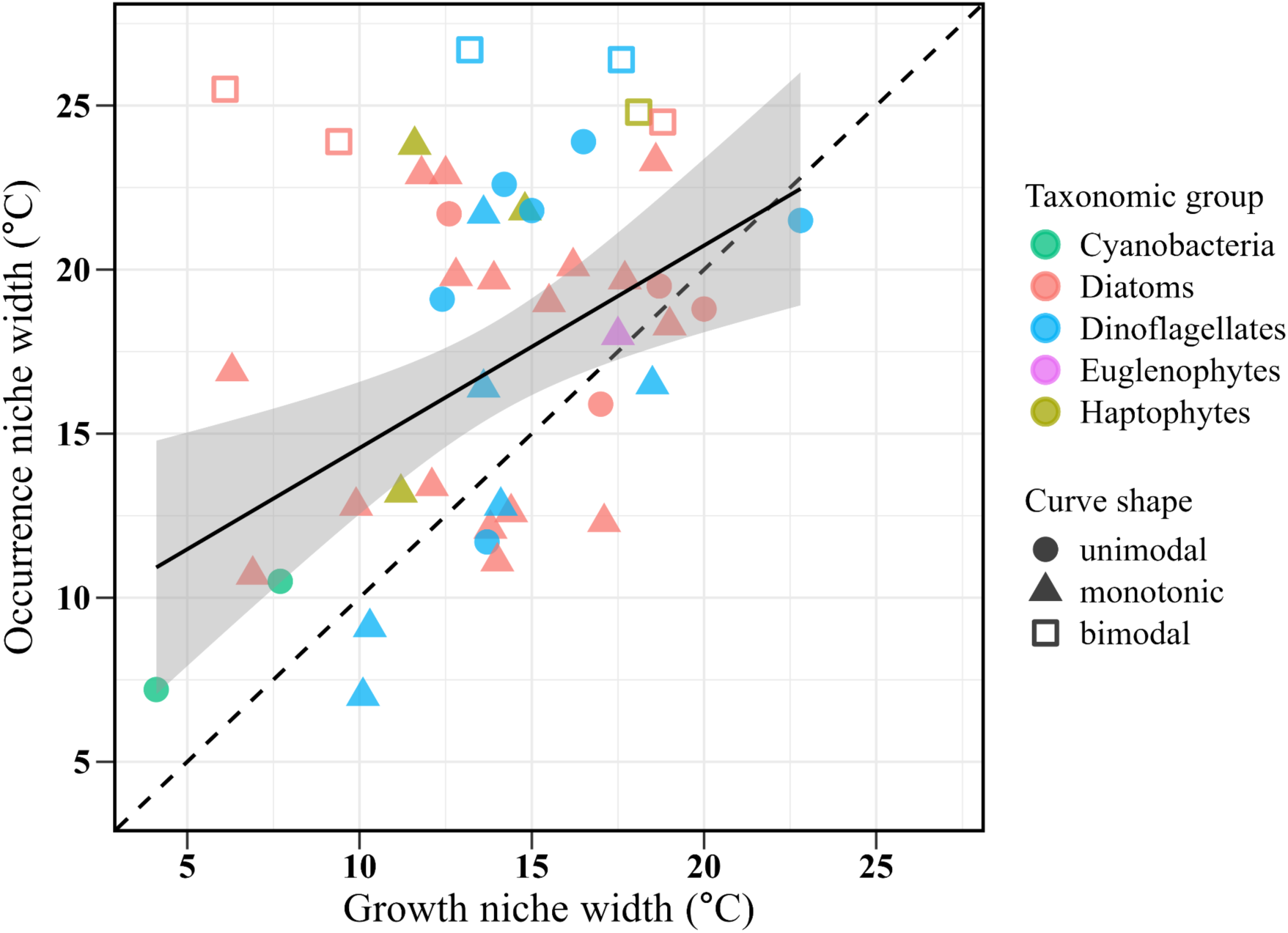
The growth niche width in the lab is positively associated with the occurrence niche width in the field (slope = 0.62, intercept = 8.4, *p* = 0.002, R^2^ = 0.24, Table S4). The residual distribution exhibits mild deviation from normality but robust regression returns nearly identical coefficients and standard errors, so we present the OLS results here. Occurrence niche width estimates are likely most reliable for unimodal curves, followed by monotonic and finally bimodal curves. The six bimodal curves (hollow squares) were therefore excluded from the regression. Where lab growth curves exist for multiple strains of the same species, points represent the niche width of the envelope of all strains’ growth curves.

## Discussion

Our analyses show that simple lab measurements can predict global species occurrence patterns well across biogeochemically-important functional groups, despite multiple sources of uncertainty and bias in both lab and field parameter estimates. In the lab data, there is substantial variation in methodology between the original studies, intraspecific variation in growth curves that we only capture partially (we have growth curves for >1 strain for only 18 out of 45 species, Fig. S1), bias towards strains that grow fast in standard lab media, local adaptation, adaptation to lab conditions (Listmann et al., 2016; O’Donnell et al., 2018; Schaum et al., 2018), and interactions with other environmental drivers that can shift parameters of temperature curves (Thomas et al., 2017; Litchman and Thomas, 2023). In the field data and associated SDMs, there are small sample sizes, spatial sampling biases, taxonomic uncertainties and biases, noise introduced by the flexible modelling approach (Merow et al., 2014), covariation and interactions between environmental predictors, as well as mismatches between the spatial scales of observations and those of the environmental predictors (Elith and Leathwick, 2009; Phillips et al., 2009; Faust and Raes, 2012; Brun et al., 2015, 2016; Leles et al., 2019). Despite these factors, the median growth temperature was reasonably strongly associated with the median occurrence temperature, and their mean relationship was close to 1:1 (Fig. 2). The association between growth and occurrence niche widths was also positive, though weaker (Fig. 4). In short, fundamental thermal niches do predict realised niches reasonably well, both in terms of central tendency and dispersion. Overall, this suggests that SDM-predicted changes in occurrence probability (or habitat suitability) could reasonably be interpreted ecologically as changes in potential growth rate. This would imply that the broad increase in phytoplankton habitat suitability across the oceans predicted in Benedetti et al. (2021a) indicates an increase in favourable growth conditions - but not necessarily higher cell concentrations - for the focal species. It also suggests that we should be able to reliably predict phytoplankton species ranges today and to forecast warming-driven shifts in the future.

These results should increase our confidence both in the utility of simple lab experiments in assessing responses to the abiotic environment (Kearney and Porter, 2009), and in the potential to infer environmental preferences from occurrence data using SDMs (Guisan et al., 2017; Benedetti et al., 2021a). It indicates that temperature is likely such a strong driver of marine microbial species distributions and ecological dynamics in nature that even relationships extracted based on limited data matched with environmental predictors at coarse spatiotemporal scales offers useful predictive power. Despite substantial evidence that interactions between environmental drivers shapes growth and thermal niches (Thomas et al., 2017; Litchman and Thomas, 2023), ignoring these interactions has not eliminated the correspondence between fundamental and realised niches in our data. This suggests that at least at the coarse scales we examine here, interactions may have a limited role in shaping presence/absence patterns, although abundance is very likely strongly influenced by them. Our findings imply that global-scale SDMs - originally developed and successfully used for terrestrial taxa (Guisan and Zimmermann, 2000) - work fairly well for marine microbes, despite reasonable prior scepticism (Dutkiewicz et al., 2020; Henson et al., 2021). It also suggests that the responses of plankton species or functional groups to changes in environmental conditions is likely to be captured in mechanistic marine ecosystem models, which rely on the parameterisation of biological agents based on theory and lab experiments (e.g., Le Quéré et al., 2005; Anderson et al., 2021).

A subtle implication is that evolution on short temporal and spatial scales (as seen in experiments; e.g., O’Donnell et al., 2018) likely does not strongly affect intraspecific variation in the temperature responses of marine phytoplankton, although it undoubtedly shapes them on longer time scales (Thomas et al., 2016). If adaptation was rapid, growth curves would have shifted substantially from their original shapes and lost their correspondence with occurrence patterns, and if local adaptation was strong, spatial variation in ocean temperatures would have similarly erased the correspondence with growth curves. Consistent with this, temperature responses do appear to be conserved to a moderate degree in ectotherms (Araujo et al., 2013). Although phytoplankton do adapt rapidly in lab experiments (Collins and Bell, 2004; Schlüter et al., 2016) including to temperature changes (Listmann et al., 2016; O’Donnell et al., 2018), the magnitude of changes observed in these studies has not been very large.

Our analyses have some important caveats arising from data limitations and modelling choices:

(i) Although the relationship between median growth and occurrence temperatures is reasonably strong, it is clearly shaped by the relatively small number of isolates collected from polar and tropical regions in our dataset. There is substantial variation in median occurrence temperatures at intermediate values of median growth temperature (approximately 15-20 ℃). This may be because the majority of the occurrence probability curves are monotonic, or because there is greater temperature variation at these latitudes. The latter may also be associated with greater unmeasured intraspecific variation and a larger mismatch between our coarse temperature grid (1°×1°, mean over monthly mixed layer depth) and the actual temperatures experienced *in situ*.
(ii) The probability of occurrence has an upper bound of 1 at all temperatures while the upper limit of growth rate increases exponentially with temperature (the Eppley curve; Eppley, 1972). We do not see a way to normalise the growth curves to obey a consistent upper bound that does not introduce additional problems for interpretation. A mathematical consequence of the difference between these two curve shapes is that median growth temperatures may be expected to be higher than the median occurrence temperatures on average. However, we see little evidence for this in our analyses.
(iii) The relatively flat occurrence probability curves (Fig. S2) suggests that many species occur across the entire marine surface temperature range of >30 ℃. In contrast, most growth curves exhibit positive growth over a range of only ∼20 ℃ in the lab, although this is at the strain and not species level (Thomas et al., 2016). The flatter occurrence probability curves and wider occurrence niches in SDM models likely reflect a mix of biological and other causes. Biologically, intraspecific variation or local adaptation should lead to occurrence niche widths being wider than growth niche widths, and many species do appear to have recorded presences across a wide temperature range (Fig. S2). However, some of these locations may represent ecological sinks, where advection has moved individuals into regions where they experience negative local population growth rates (Pulliam, 2000). Data limitations such as the coarse spatiotemporal scale of the environmental data, misidentification, patchy sampling, and covariance of temperature with other environmental drivers also shape this pattern. Finally, algorithms such as GLMs and GAMs will smoothen sharp response curves, possibly leading to flatter and broader responses than the parametric TPC equation used for the growth curves. For these reasons, we chose not to investigate thermal limits (*T*_max_ and *T*_min_) or quantile-based proxies for them, despite their value in predicting future species distributions.
(iv) In cases where we have multiple measured growth curves for a species, the intraspecific variation affects our estimates of growth niche width in particular (Fig. S1). Incorporation of this intraspecific variation via the envelope approach we use strengthens the niche width relationship slightly (R^2^ = 0.24 vs. 0.18; compare Figs. 4 & S9); in contrast, the median growth temperatures are relatively insensitive to this (compare Figs. 2 & S6). This indicates that growth niche width estimates for species with just one strain’s growth curve are underestimated and is consistent with the observed pattern of occurrence niche widths being wider than growth niche widths in general (Fig. 4). Measuring additional strains of each species could therefore strengthen this relationship (Smith et al., 2021; Krinos et al., 2025).
(v) It is difficult to believe that bimodal occurrence probability curves accurately capture underlying ecophysiology and so we excluded them from our analyses. Bimodality may reflect the existence of cryptic species (Bickford et al., 2007), the effect of biotic interactions (Lalli and Parsons, 1997; Worden et al., 2015), spatial biases in sampling and pseudoabsence generation (Phillips et al., 2009), or environmental covariation (Dormann et al., 2013), exacerbated by our use of flexible modelling approaches (neural networks). The authors have found in previous work on plankton SDMs that species such as *E. huxleyi* that exhibit boom-and-bust dynamics are difficult to constrain, as their dynamics are likely heavily driven by changes in nutrient concentrations at spatiotemporal scales that we do not have environmental data for (weekly, local scales, high patchiness). In the case of *E. huxleyi*, the existence of multiple attested ecotypes that are not well-sampled may contribute to its bimodality. The strong governing role of nutrients likely confounds the apparent temperature response, especially because we do not account for the complex temperature-nutrient interaction in the models.
(vi) Our data represents between 10% and 97% of the occurrences in the Phytobase v1 dataset, depending on the functional group. While this is not a trivial fraction, it is also not a random sample, being biased towards species that are easier to recognise and easy to grow in the lab. And these datasets rely on morphological identification, while genetics-based species diversity estimates tend to be orders of magnitude larger (Vincent et al., 2022). Therefore, a great many taxa are excluded, including *Prochlorococcus* sp. and *Synechococcus* sp., which form the majority of cyanobacterial records but are not identified to a species level. Others are conflated, and so future work using genetics-based SDMs may find that occurrence niche widths are narrower than those we estimate here.

We consider this to be a step towards linking ecophysiology, occurrences, and ecological forecasts, and foresee multiple ways in which this can be built upon. On the experimental front, we recommend experimentally measuring growth curves of additional species, especially those that are biogeochemically or ecologically important. New strains of already-measured species from different environments would be useful as well for improving growth niche width estimates. It would likely also help to measure interactions between temperature and other drivers (Thomas et al., 2017; Seifert et al., 2020; Kremer et al., 2025), taking advantage of efficient experimental designs (Collins et al., 2022; Thomas and Ranjan, 2024), as these interactions likely determine distributions and the bimodal curves we see for some species (especially boom-and-bust species). On the occurrence data front, our analyses suggest that for most species, it would be most valuable to increase the number of presences/pseudoabsences in the hottest and coldest waters (although for the bimodal curves, data from intermediate temperatures may be more useful). The presence/pseudoabsence data limitations we face here may be ameliorated substantially by using new flow cytometry (Ricour, 2023) and omics data sources such as the Tara Oceans project (de Vargas et al., 2015; Sunagawa et al., 2020; https://end.mio.osupytheas.fr/Ecological_Niche_database/).

On the SDM front, there are multiple avenues to build on. One promising step would involve imposing constraints such as unimodality or monotonicity on flexible models such as GAMs (Pya and Wood, 2015; Citores et al., 2020, Valle et al., 2023). An alternative would involve using parametric models of temperature-dependent presence/absence, perhaps with the same broad shape as the functions used to describe growth curves. This mechanistic species distribution modelling or mechanistic niche modelling (Kearney and Porter, 2009) approach has had recent success in explaining fish distributions at a small geographical scale (Wagner et al., 2023), and the estimated parameters can be compared directly with those from growth curves. They can also incorporate the dependence of growth on multiple environmental dimensions and compare the resulting biogeographic patterns with observed or inferred species ranges from presence/pseudoabsence data (Seifert et al., 2023). Accounting for the dependence of median temperature and occurrence niche width on additional drivers may strengthen the relationships we observe here, resolve some of the anomalous bimodal curves, and perhaps explain the latitudinal pattern in the difference between the median growth and occurrence temperatures (Fig. 3). A third approach would involve evaluating correspondence between entire growth and occurrence probability curves (perhaps using functional data analysis), not just the simple measures of central tendency and dispersion that we compare here. And finally, SDMs modelling biomass or abundance rather than occurrence are likely to be both more accurate and useful. Preliminary studies have found good correspondence between areas with high species-specific habitat suitability (identical to areas of high occurrence probability) and areas with high species-specific biomass, for a range of plankton species. This suggests that biomass-based estimates of niche centres and widths are within reach (e.g., Schickele et al., 2025). In any case, we believe it would be worthwhile to identical and link meaningful temperature limits (*T*_max_ and *T*_min_) in growth and occurrence probability, and to model interactions between temperature and other drivers (Geider et al., 1998; Ross and Geider, 2009; Thomas et al., 2017; Litchman and Thomas, 2023; Kremer et al., 2025).

Perhaps the most useful additional direction would be to apply our approach at the level of functional groups/plankton functional types (PFTs) instead of species. These PFTs are represented in ocean biogeochemical and ecosystem models and are therefore at the core of oceanographers’ efforts to forecast changes to future oceans and climate (e.g., Le Quéré et al., 2005; Follows et al., 2007; Dutkiewicz et al., 2020). At present, obtaining useful forecasts from these models requires extensive ‘tuning’ of multiple unknown parameters, which in turn depends on modellers’ knowledge of the present-day distributions of different PFTs and their underlying causes. It would be straightforward to estimate the parameters of occurrence and growth curves at the PFT level to evaluate SDM accuracy. This could be performed for temperature and other drivers as well. The envelope-based approach may need revisiting for nutrients and light, and possibly the splitting of existing PFTs into subgroups with similar ecological niches (Benedetti et al., 2018, 2023b). Agreement between growth curves and SDMs at the PFT level would provide a strong grounding for biogeochemical and ecosystem models, and perhaps allow some of their free parameters to be constrained better. This could in turn lead to better predictions of future productivity and biogeochemistry.

## Conclusion

Our work shows that in phytoplankton, fundamental thermal niches in lab experiments predict realised thermal niches reasonably well. Given that data availability and methods will only improve, this is cause for optimism in the effort to forecast important marine ecosystem properties: species distributions, plankton biogeography, community composition, biodiversity patterns, and ecosystem dynamics. Furthermore, since temperature is reliably diagnosed from satellite measurements, we may be able to develop reliable real-time forecasts of plankton community composition using remote sensing, at least in areas where temperature is the dominant driver of plankton biogeography. As phytoplankton form the foundation of aquatic ecosystems and monitoring their populations in the field is expensive, validating links between fundamental niches, realised niches, and model-predicted community composition would be an efficient way to improve predictions of ecosystem processes, productivity and biodiversity. We therefore believe that this provides a strong argument for increasing experimental measurements of ecophysiology with the explicit goal of making and testing predictions across space and time, linking experiments with the ever-increasing observation datasets and vital biogeochemical and ecosystem modelling efforts.

## Acknowledgements

We thank Colin T. Kremer & Maya Chari for providing us with TPC parameter estimates for the growth curves of 17 strains belonging to 6 species. The work presented in this article results in part from funding provided by national committees of the Scientific Committee on Oceanic Research (SCOR) and from a grant to SCOR from the US National Science Foundation (OCE-1840868) to the Changing Oceans Biological Systems project. T.L. acknowledges funding from the National Natural Science Foundation of China (No. 42506141) and Shandong Provincial Natural Science Foundation (No. ZR2025QC1894Z). F.B. and M.V. acknowledge funding from the European Union’s Horizon 2020 Research and Innovation Programme under grant agreement no. 862923 (AtlantECO) and under grant agreement no. 101059915 (BIOceans5D). This output reflects only the authors’ views, and the European Union cannot be held responsible for any use that may be made of the information contained therein.

## Author contributions

**Ting Lv**: conceptualization, data curation, formal analysis, funding acquisition, investigation, methodology, software, validation, visualization, writing – original draft, writing – review and editing. **Fabio Benedetti**: conceptualization, data curation, formal analysis, methodology, software, validation, writing – review and editing. **Dominic Eriksson**: data curation, formal analysis, methodology, validation, writing – review and editing. **Meike Vogt**: conceptualization, data curation, funding acquisition, methodology, validation, writing – review and editing. **Mridul K. Thomas**: conceptualization, data curation, formal analysis, investigation, methodology, project administration, supervision, validation, visualization, writing – original draft, writing – review and editing.

