## Supplementary material for "Phytoplankton performance in the lab predicts occurrence in the field across a global temperature gradient": S1; S2; S3; S4; S5; S6; S7; S8; S9; S10

#### Supplementary tables

**Table S1**. The number of occurrences of each of the 45 species in our field data, along with metrics of model skill for the SDM ensemble. The Area Under the Receiver Operating Characteristic curve (AUC) ranges between 0 and 1, with values <0.5 indicating worse than random model skill (Guisan et al., 2017). The True Skill Statistic (TSS) values range between −1 and +1, with null values indicating that models perform no better than at random (Allouche et al., 2006).

| Taxonomic group | Species | Number of occurrences | Number of pseudo-absences | Mean AUC | Stdev AUC | Mean TSS | Stdev TSS |
|---|---|---|---|---|---|---|---|
| Diatoms | *Asterionellopsis glacialis* | 1026 | 10260 | 0.728 | 0.026 | 0.338 | 0.040 |
| | *Chaetoceros affinis* | 1077 | 10769 | 0.878 | 0.011 | 0.667 | 0.025 |
| | *Chaetoceros didymus* | 838 | 8378 | 0.816 | 0.019 | 0.487 | 0.031 |
| | *Chaetoceros lorenzianus* | 1010 | 10101 | 0.924 | 0.009 | 0.750 | 0.020 |
| | *Chaetoceros pseudocurvisetus* | 239 | 2390 | 0.818 | 0.021 | 0.474 | 0.074 |
| | *Chaetoceros simplex* | 102 | 1020 | 0.847 | 0.024 | 0.578 | 0.066 |
| | *Corethron pennatum* | 2479 | 24790 | 0.757 | 0.010 | 0.388 | 0.020 |
| | *Coscinodiscus concinnus* | 879 | 8790 | 0.793 | 0.012 | 0.447 | 0.020 |
| | *Coscinodiscus granii* | 420 | 4201 | 0.876 | 0.018 | 0.640 | 0.039 |
| | *Coscinodiscus wailesii* | 399 | 3992 | 0.833 | 0.024 | 0.515 | 0.052 |
| | *Dactyliosolen fragilissimus* | 1140 | 11400 | 0.722 | 0.017 | 0.329 | 0.031 |
| | *Detonula confervacea* | 487 | 4870 | 0.808 | 0.018 | 0.506 | 0.036 |
| | *Ditylum brightwellii* | 1132 | 11319 | 0.782 | 0.019 | 0.431 | 0.034 |
| | *Eucampia zodiacus* | 816 | 8161 | 0.785 | 0.024 | 0.448 | 0.048 |
| | *Fragilariopsis kerguelensis* | 161 | 1610 | 0.969 | 0.020 | 0.885 | 0.045 |
| | *Helicotheca tamesis* | 178 | 1780 | 0.876 | 0.029 | 0.627 | 0.063 |
| | *Leptocylindrus danicus* | 1307 | 13071 | 0.783 | 0.012 | 0.448 | 0.029 |
| | *Proboscia inermis* | 172 | 1722 | 0.777 | 0.039 | 0.402 | 0.091 |
| | *Rhizosolenia setigera* | 989 | 9891 | 0.803 | 0.012 | 0.479 | 0.021 |
| | *Skeletonema costatum* | 1891 | 18907 | 0.692 | 0.018 | 0.310 | 0.021 |
| | *Stephanopyxis palmeriana* | 268 | 2680 | 0.856 | 0.024 | 0.584 | 0.053 |
| | *Thalassionema nitzschioides* | 5868 | 46901 | 0.826 | 0.009 | 0.543 | 0.010 |
| | *Thalassiosira gravida* | 613 | 6135 | 0.833 | 0.024 | 0.538 | 0.042 |
| | *Thalassiosira nordenskioeldii* | 172 | 1722 | 0.872 | 0.039 | 0.646 | 0.073 |
| Haptophytes | *Calcidiscus leptoporus* | 645 | 6453 | 0.736 | 0.013 | 0.376 | 0.035 |

| | | | | | | | |
|---|---|---|---|---|---|---|---|
| | *Coccolithus pelagicus* | 603 | 6031 | 0.708 | 0.023 | 0.319 | 0.025 |
| | *Emiliania huxleyi* | 1730 | 8733 | 0.645 | 0.033 | 0.214 | 0.038 |
| | *Phaeocystis pouchetii* | 330 | 3300 | 0.948 | 0.011 | 0.814 | 0.026 |
| Cyanobacteria | *Crocosphaera watsonii* | 243 | 2434 | 0.753 | 0.027 | 0.437 | 0.047 |
| | *Trichodesmium erythraeum* | 296 | 2680 | 0.801 | 0.031 | 0.491 | 0.050 |
| Dinoflagellates | *Akashiwo sanguinea* | 269 | 2691 | 0.852 | 0.021 | 0.589 | 0.043 |
| | *Dinophysis caudata* | 958 | 9581 | 0.921 | 0.006 | 0.741 | 0.016 |
| | *Leonella granifera* | 108 | 1081 | 0.934 | 0.020 | 0.783 | 0.055 |
| | *Prorocentrum balticum* | 148 | 1479 | 0.784 | 0.070 | 0.484 | 0.112 |
| | *Prorocentrum cordatum* | 249 | 2492 | 0.676 | 0.041 | 0.234 | 0.081 |
| | *Prorocentrum gracile* | 411 | 4109 | 0.928 | 0.009 | 0.701 | 0.024 |
| | *Prorocentrum lima* | 128 | 1280 | 0.787 | 0.041 | 0.414 | 0.091 |
| | *Prorocentrum micans* | 647 | 6469 | 0.789 | 0.012 | 0.456 | 0.030 |
| | *Scrippsiella acuminata* | 478 | 4780 | 0.850 | 0.024 | 0.613 | 0.048 |
| | *Thoracosphaera heimii* | 185 | 1850 | 0.928 | 0.018 | 0.744 | 0.048 |
| | *Tripos furca* | 5253 | 11940 | 0.662 | 0.008 | 0.243 | 0.014 |
| | *Tripos fusus* | 8360 | 11940 | 0.576 | 0.010 | 0.112 | 0.014 |
| | *Tripos lineatus* | 3148 | 11940 | 0.722 | 0.015 | 0.342 | 0.021 |
| | *Tripos muelleri* | 5283 | 11940 | 0.643 | 0.011 | 0.209 | 0.023 |
| Euglenophytes | *Eutreptiella gymnastica* | 165 | 1649 | 0.855 | 0.023 | 0.536 | 0.063 |

**Table S2**. Model summary for Figure 2: the median occurrence temperature as a function of median growth temperature.

| | median occurrence temperature (°C) | | |
|---|---|---|---|
| Predictors | Estimates | CI | *p*-value |
| (Intercept) | 1.60 | -4.03 – 7.24 | 0.568 |
| median growth temperature (°C) | 0.85 | 0.56 – 1.13 | <0.001 |
| Observations | 39 | | |
| $R^2$ / $R^2$ adjusted | 0.487 / 0.473 | | |

**Table S3**. Model summary for Figure 3b: the difference between median growth and median occurrence temperatures as a function of absolute latitude.

| | Median growth temperature - Median occurrence temperature (°C) | | |
|---|---|---|---|
| Predictors | Estimates | CI | bootstrap *p*-value |
| (Intercept) | 6.50 | 4.14 – 8.87 | <0.001 |
| Absolute latitude | -0.15 | -0.19 – -0.10 | <0.001 |
| Random Effects | | | |
| $\sigma^2$ | 2.76 | | |
| $\tau_{00\ species}$ | 21.13 | | |
| ICC | 0.88 | | |
| $N_{species}$ | 35 | | |
| Observations | 65 | | |
| Marginal $R^2$ / Conditional $R^2$ | 0.179 / 0.905 | | |

**Table S4**. Model summary for Figure 4: the occurrence niche width as a function of growth niche width.

| | occurrence niche width | | |
|---|---|---|---|
| Predictors | Estimates | CI | *p*-value |
| (Intercept) | 8.39 | 3.10 – 13.68 | 0.003 |
| growth niche width | 0.62 | 0.25 – 0.98 | 0.002 |
| Observations | 39 | | |
| $R^2$ / $R^2$ adjusted | 0.239 / 0.219 | | |

### Supplementary figures

**Fig. S1**. The lab growth rate curves for all the species we study in this paper. Each black line represents a single published curve and the blue line represents the envelope for that species (i.e. the maximum growth rate achieved across all strains at any temperature). The red vertical line indicates the median growth temperature, and the shaded grey area represents the growth niche width, both calculated from the envelope.

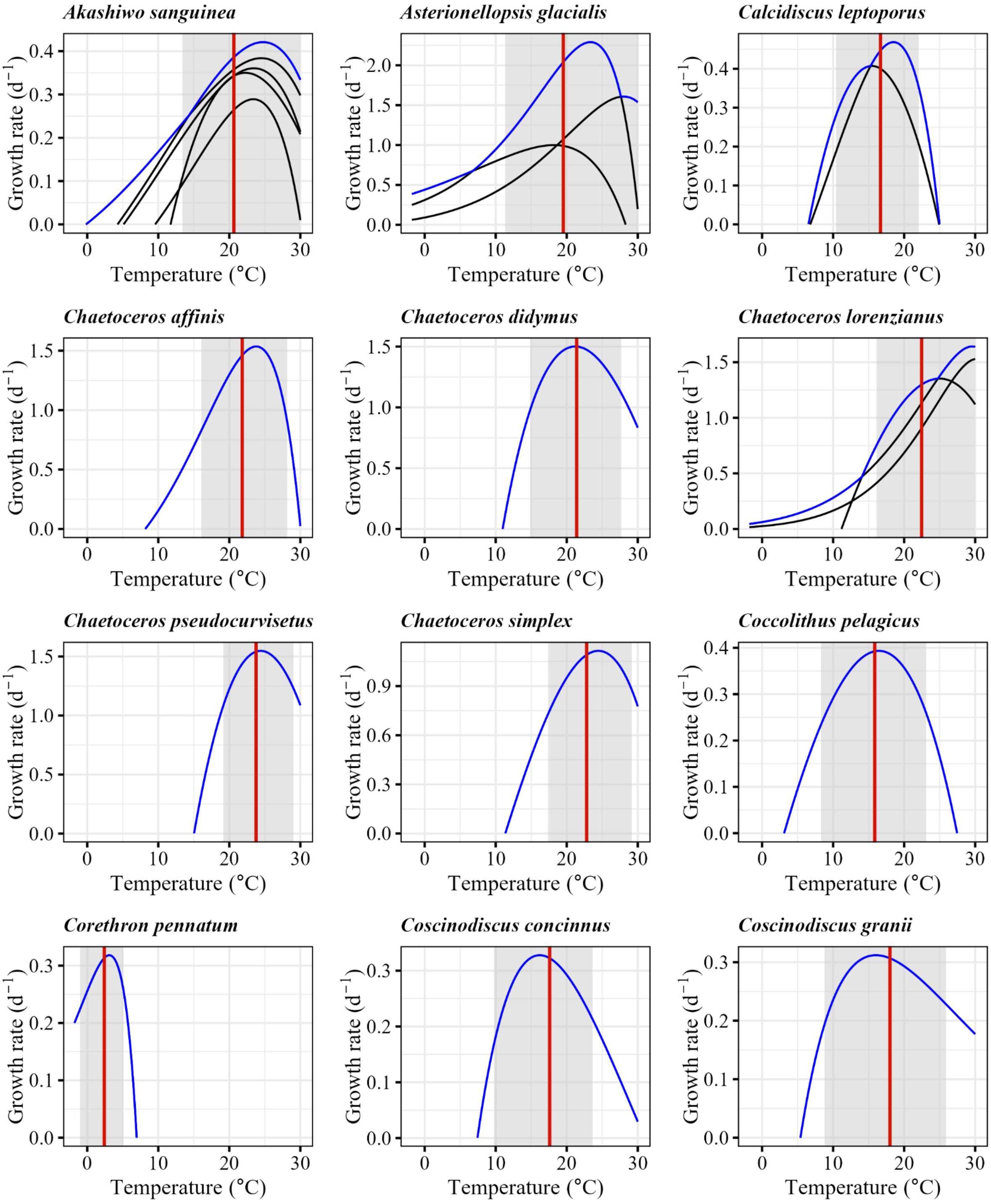

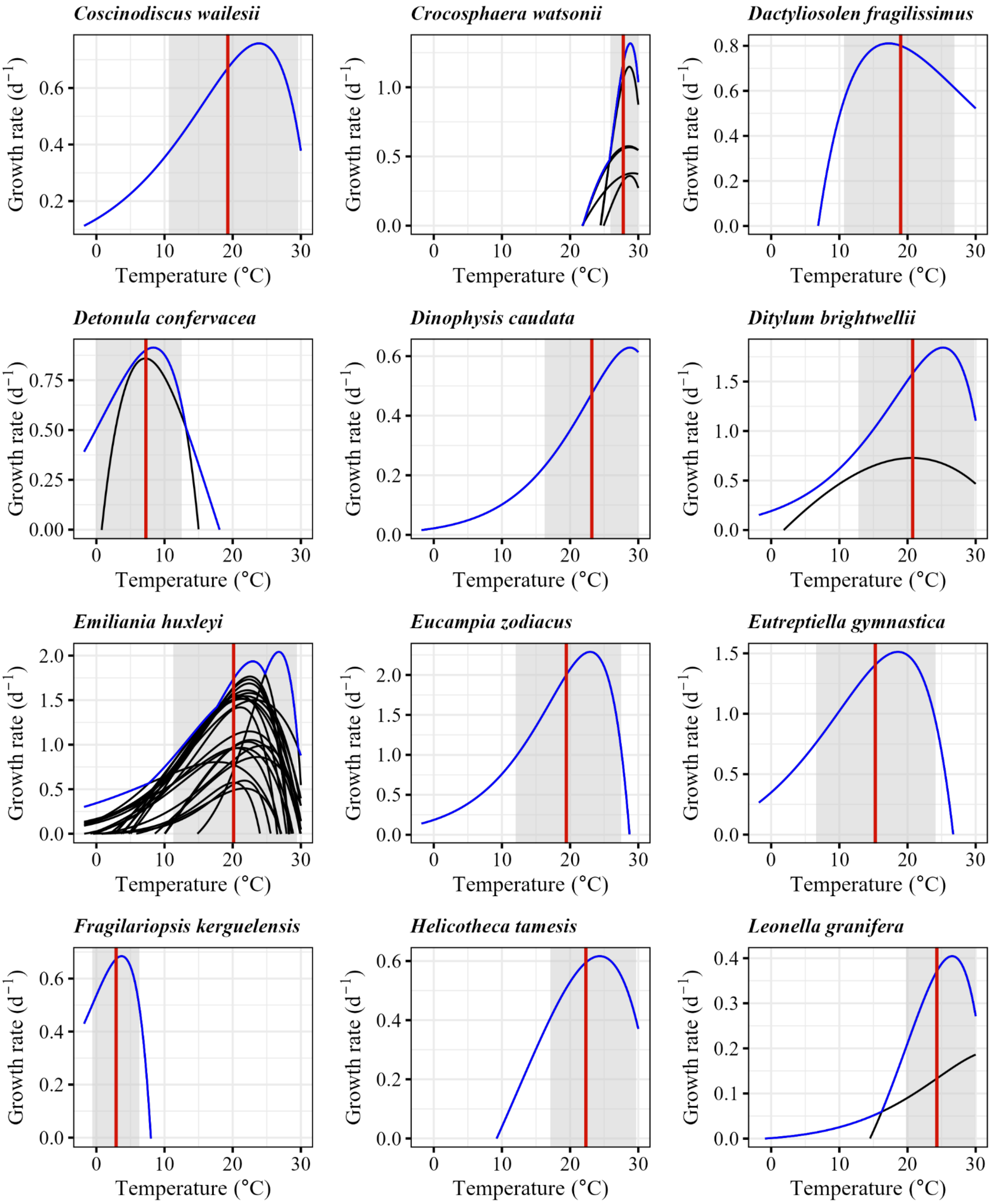
Coscinodiscus wailesii
Crocosphaera watsonii
Dactyliosolen fragilissimus
Detonula confervacea
Dinophysis caudata
Ditylum brightwellii
Emiliania huxleyi
Eucampia zodiacus
Eutreptiella gymnastica
Fragilariopsis kerguelensis
Helicotheca tamesis
Leonella granifera
Growth rate (d$^{-1}$)
Temperature (°C)

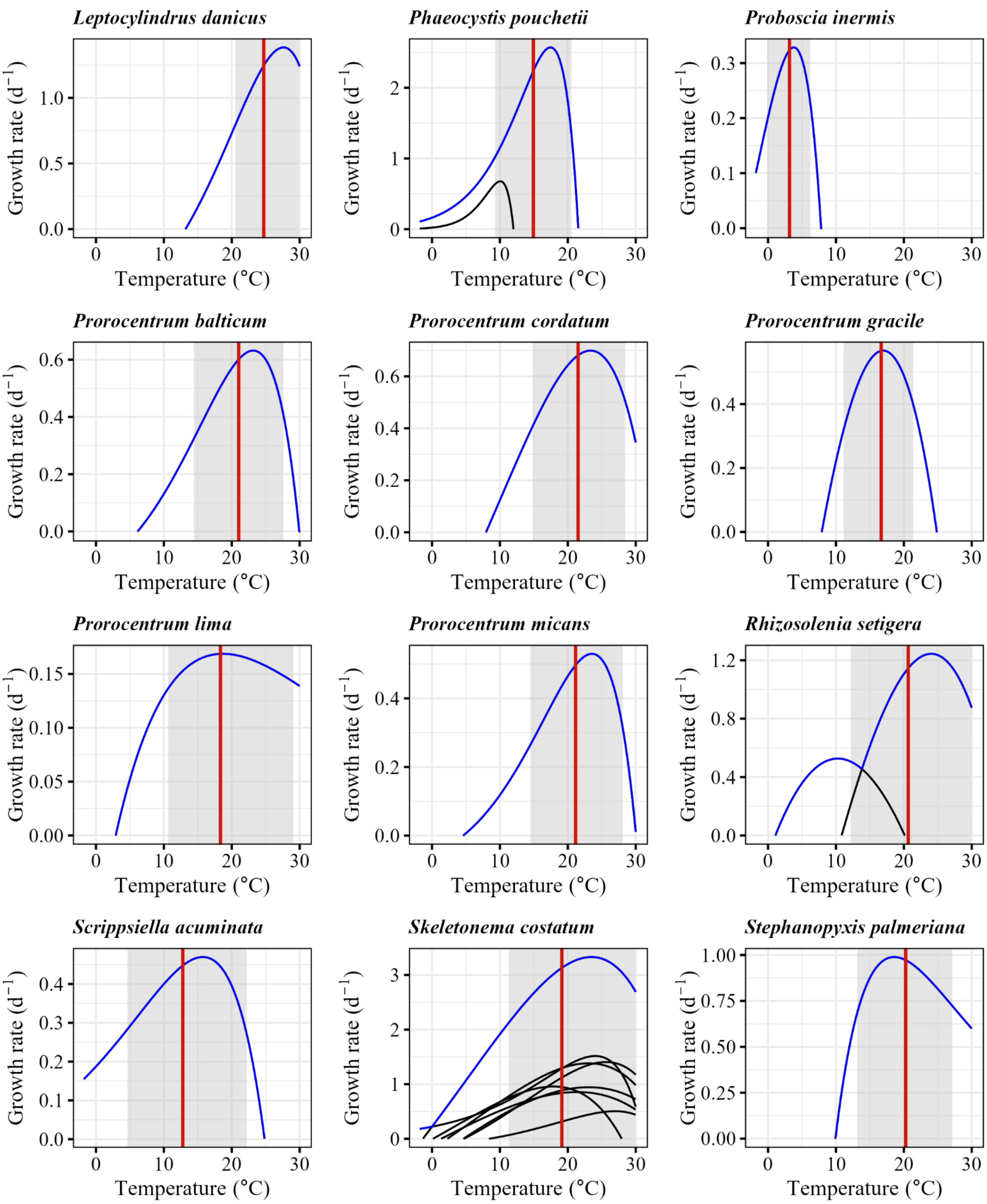
Leptocylindrus danicus
Phaeocystis pouchetii
Proboscia inermis
Prorocentrum balticum
Prorocentrum cordatum
Prorocentrum gracile
Prorocentrum lima
Prorocentrum micans
Rhizosolenia setigera
Scrippsiella acuminata
Skeletonema costatum
Stephanopyxis palmeriana
Growth rate (d$^{-1}$)
Temperature (°C)

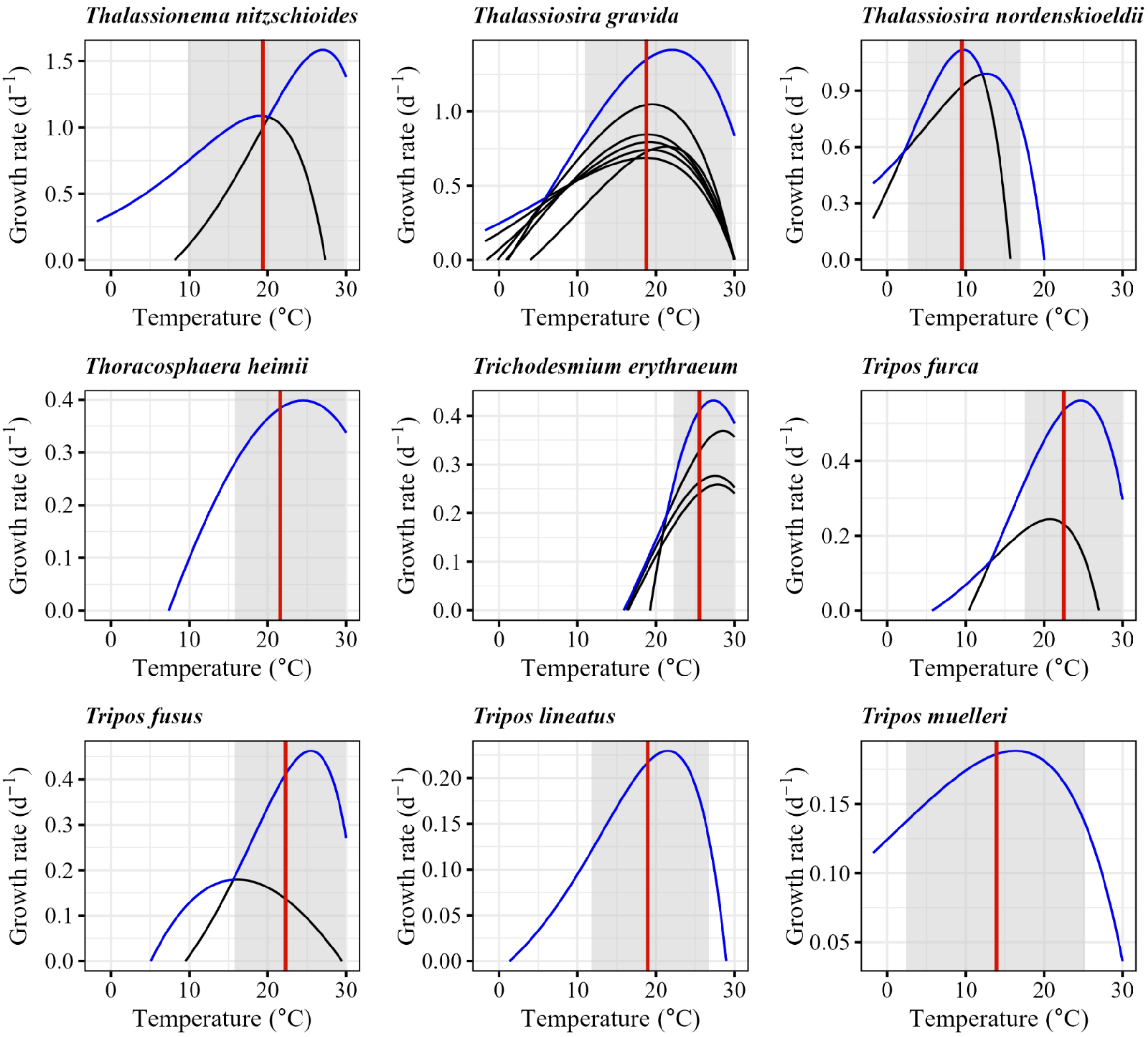

Thalassionema nitzschioides
Thalassiosira gravida
Thalassiosira nordenskioeldii
Thoracosphaera heimii
Trichodesmium erythraeum
Tripos furca
Tripos fusus
Tripos lineatus
Tripos muelleri
Growth rate (d⁻¹)
Temperature (°C)

**Fig. S2**. The field occurrence probability curves for all the species we study in this paper. The red vertical line indicates the median occurrence temperature, and the shaded gray area represents the occurrence niche width.

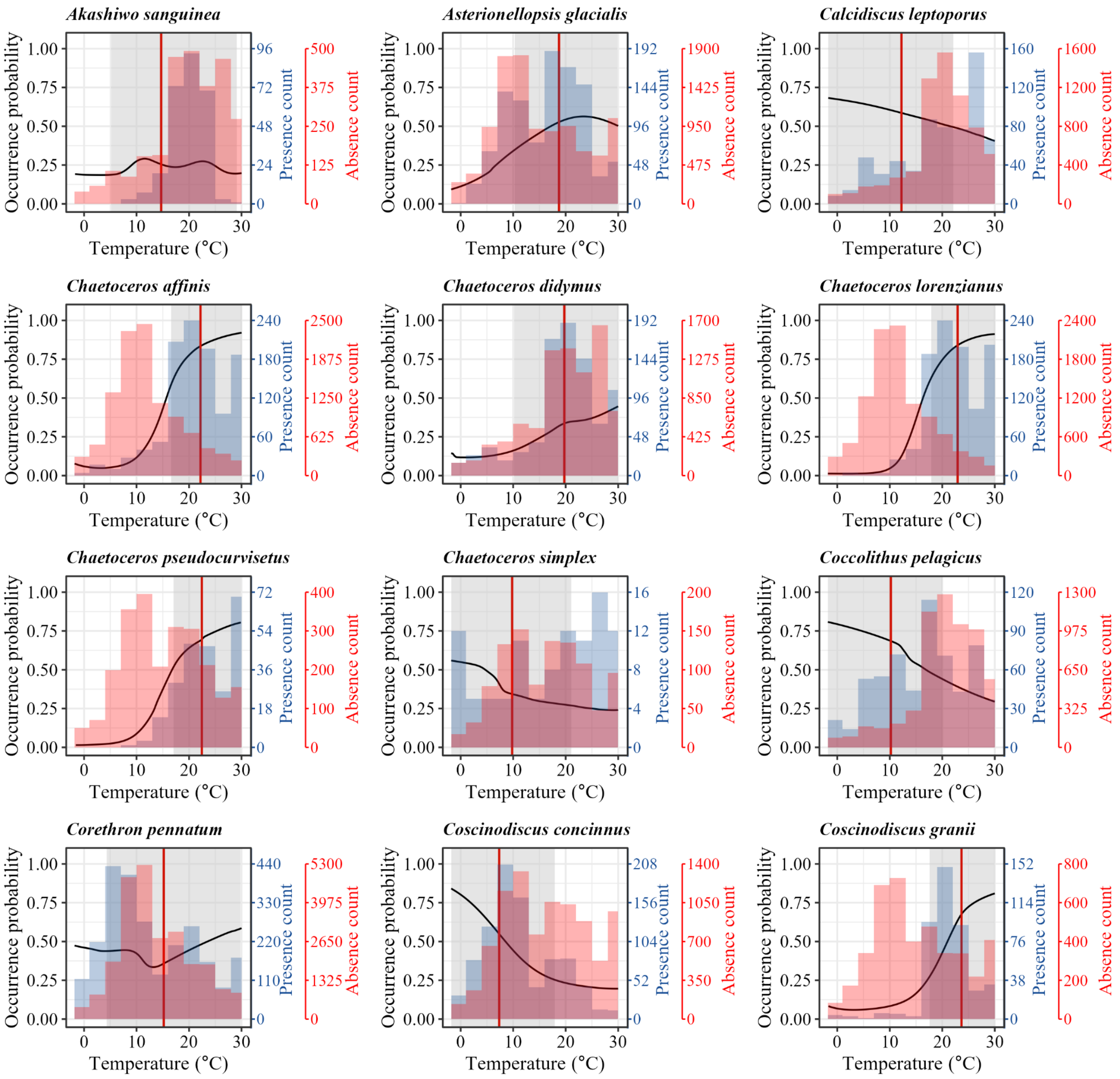

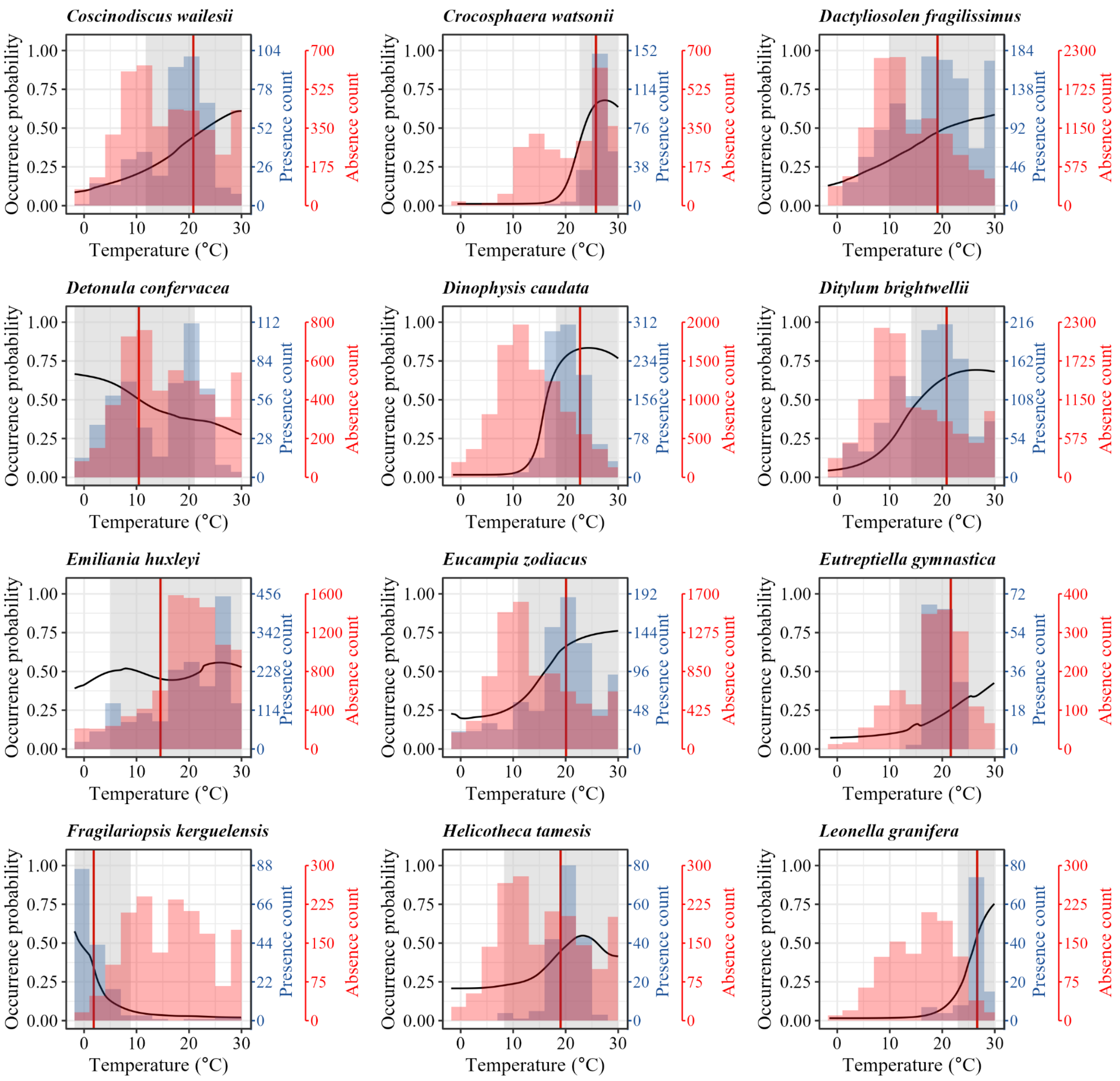
Coscinodiscus wailesii
Crocosphaera watsonii
Dactyliosolen fragilissimus
Detonula confervacea
Dinophysis caudata
Ditylum brightwellii
Emiliania huxleyi
Eucampia zodiacus
Eutreptiella gymnastica
Fragilariopsis kerguelensis
Helicotheca tamesis
Leonella granifera
Occurrence probability
Presence count
Absence count
Temperature (°C)

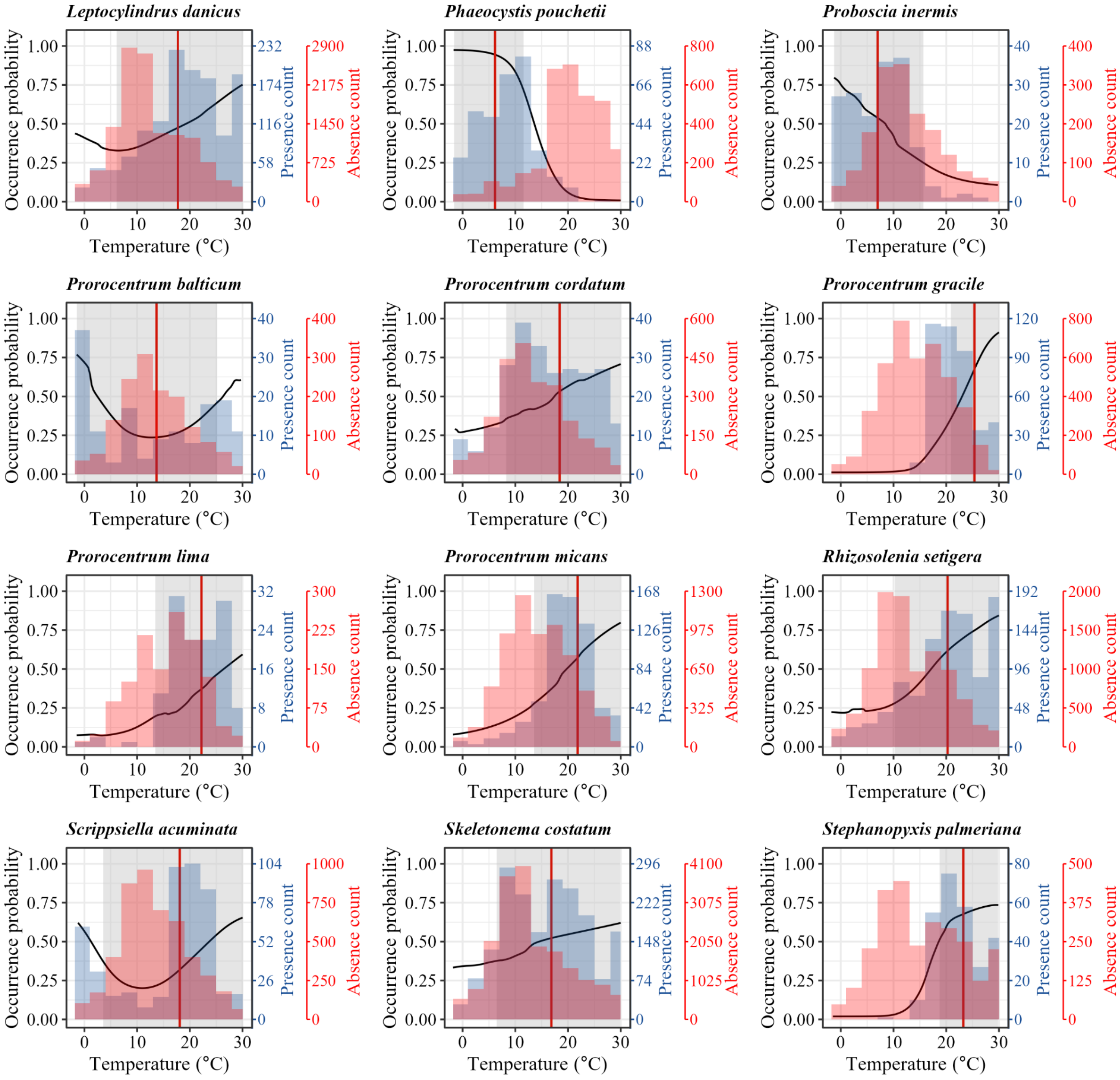
Leptocylindrus danicus
Phaeocystis pouchetii
Proboscia inermis
Prorocentrum balticum
Prorocentrum cordatum
Prorocentrum gracile
Prorocentrum lima
Prorocentrum micans
Rhizosolenia setigera
Scrippsiella acuminata
Skeletonema costatum
Stephanopyxis palmeriana
Occurrence probability
Presence count
Absence count
Temperature (°C)

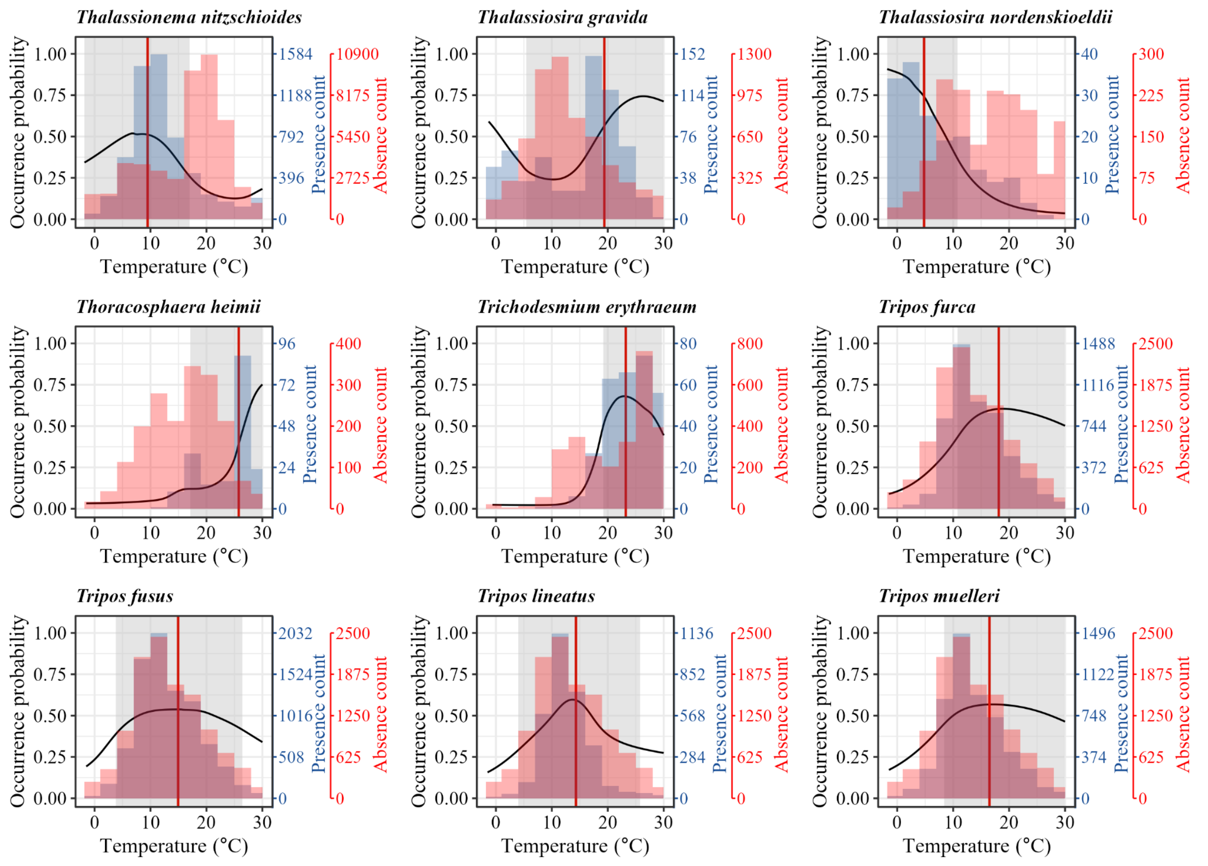
Thalassionema nitzschioides
Thalassiosira gravida
Thalassiosira nordenskioeldii
Thoracosphaera heimii
Trichodesmium erythraeum
Tripos furca
Tripos fusus
Tripos lineatus
Tripos muelleri
Occurrence probability
Presence count
Absence count
Temperature (°C)
0.00
0.25
0.50
0.75
1.00
0
10
20
30
1584
1188
792
396
10900
8175
5450
2725
152
114
76
38
1300
975
650
325
40
30
20
10
300
225
150
75
96
72
48
24
400
300
200
100
80
60
40
20
800
600
400
200
1488
1116
744
372
2500
1875
1250
625
2032
1524
1016
508
1136
852
568
284
1496
1122
748
374

**Fig. S3**. Relationships between growth niche width and occurrence niche width estimated using different cumulative area thresholds (40%, 50%, 60%, 70%, 80%, and 90% of the total area under the curve). The six species with bimodal occurrence curves (hollow squares) were excluded from the regression, as we consider their estimates unreliable. The solid lines are the linear regressions and the dashed lines are 1:1 lines. The results were qualitatively consistent across thresholds, and the 70% and 80% thresholds yielded the highest $R^2$ values.

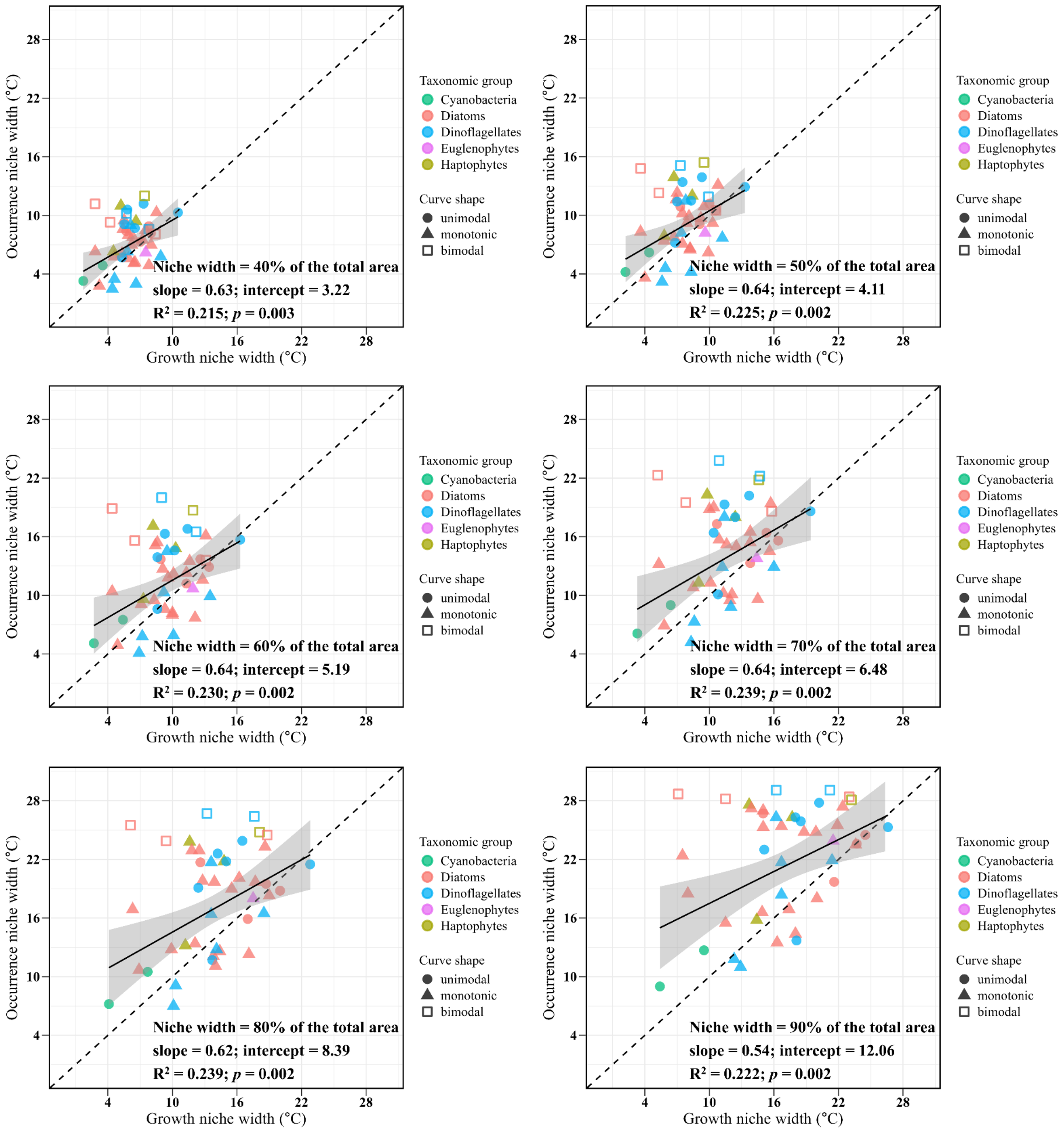

**Fig. S4**. The temperature of maximum growth rate (also known as the optimum temperature for growth) estimated from lab growth curves is very similar to the median growth temperature metric we use in our study. The dashed line represents the 1:1 line. Each point was calculated based on the envelope of all strains' growth curves.

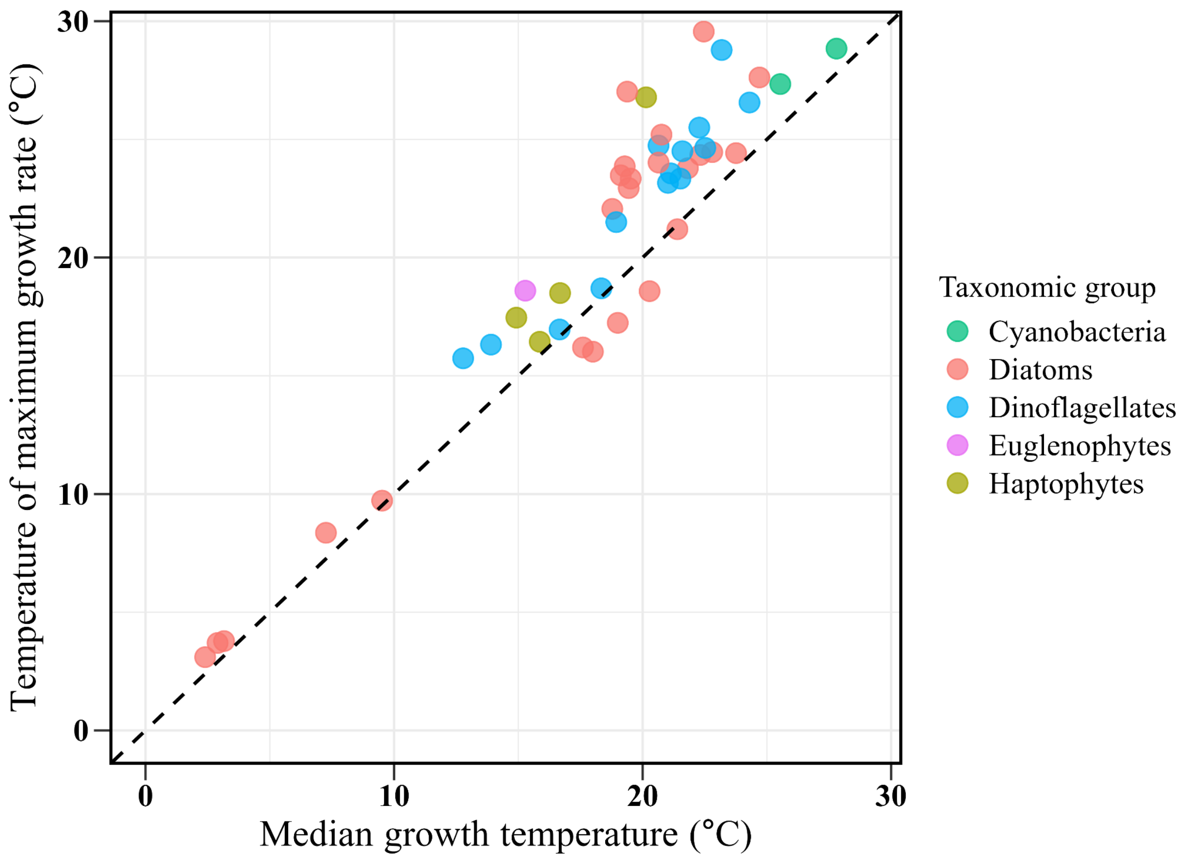

**Fig. S5**. Field-based estimates of optimum temperature (the temperature at which estimated probability of occurrence is highest) are strongly but nonlinearly associated with median occurrence temperature (unlike the linear, almost 1:1 relationship in the analogous parameters in lab growth curves; Fig. S4). This sigmoidal relationship is because the majority of the field occurrence probability curves are monotonic, leading to a bimodal distribution of 'maximum occurrence probability temperature' (the subset of unimodal curves exhibits the expected linear near 1:1 relationship). This bimodal distribution captures little relevant ecophysiological variation. This is why we do not use this measure of field temperature preference, and instead use the median occurrence temperature, which has a more continuous distribution. We believe that a greater proportion of occurrence probability curves would be found to be unimodal with better sampling of the tropics and poles. Once this data exists, it may be worth switching to the use of maximum occurrence probability temperature (and optimum temperature for growth in the case of growth curves).

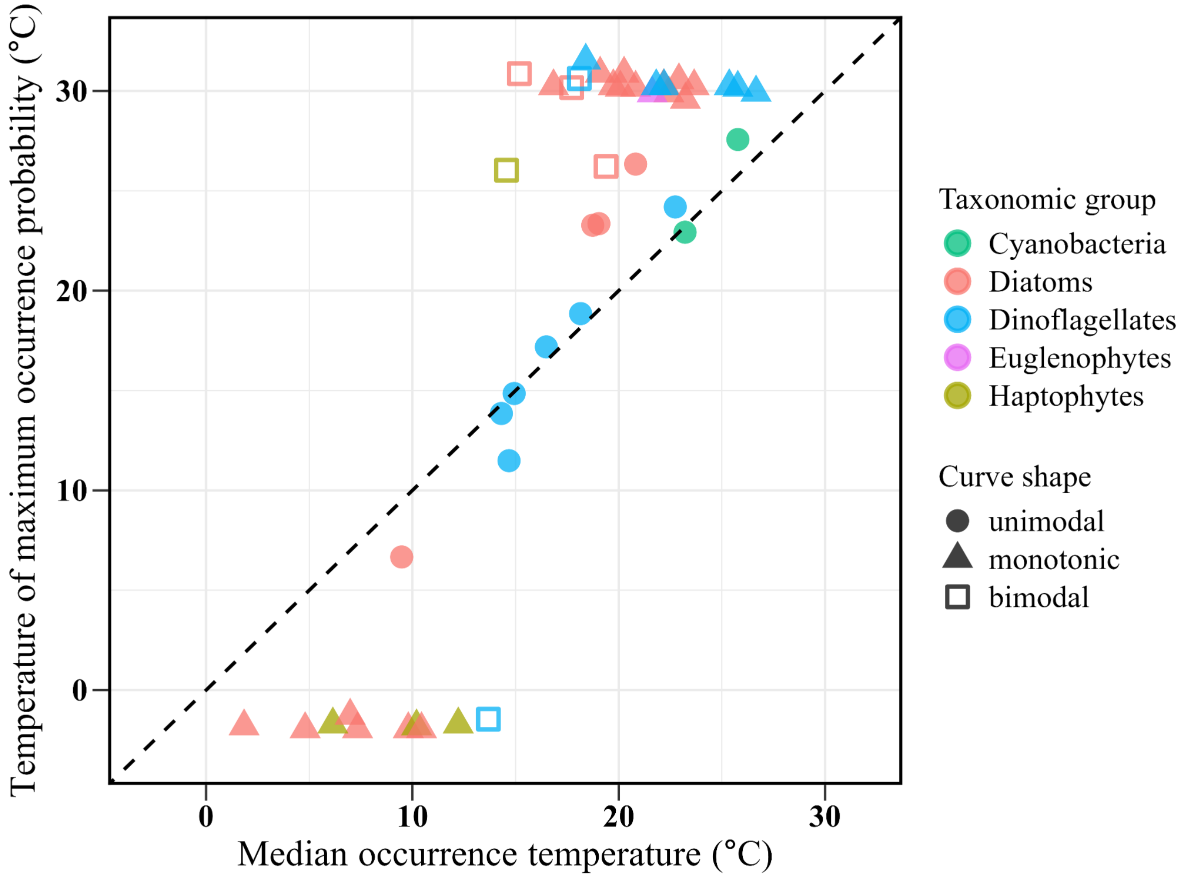

**Fig. S6**. Deviations from the relationship between median growth temperature and median occurrence temperature do not appear to be related to SDM quality (as indicated by AUC and TSS) or with the number of occurrence records.

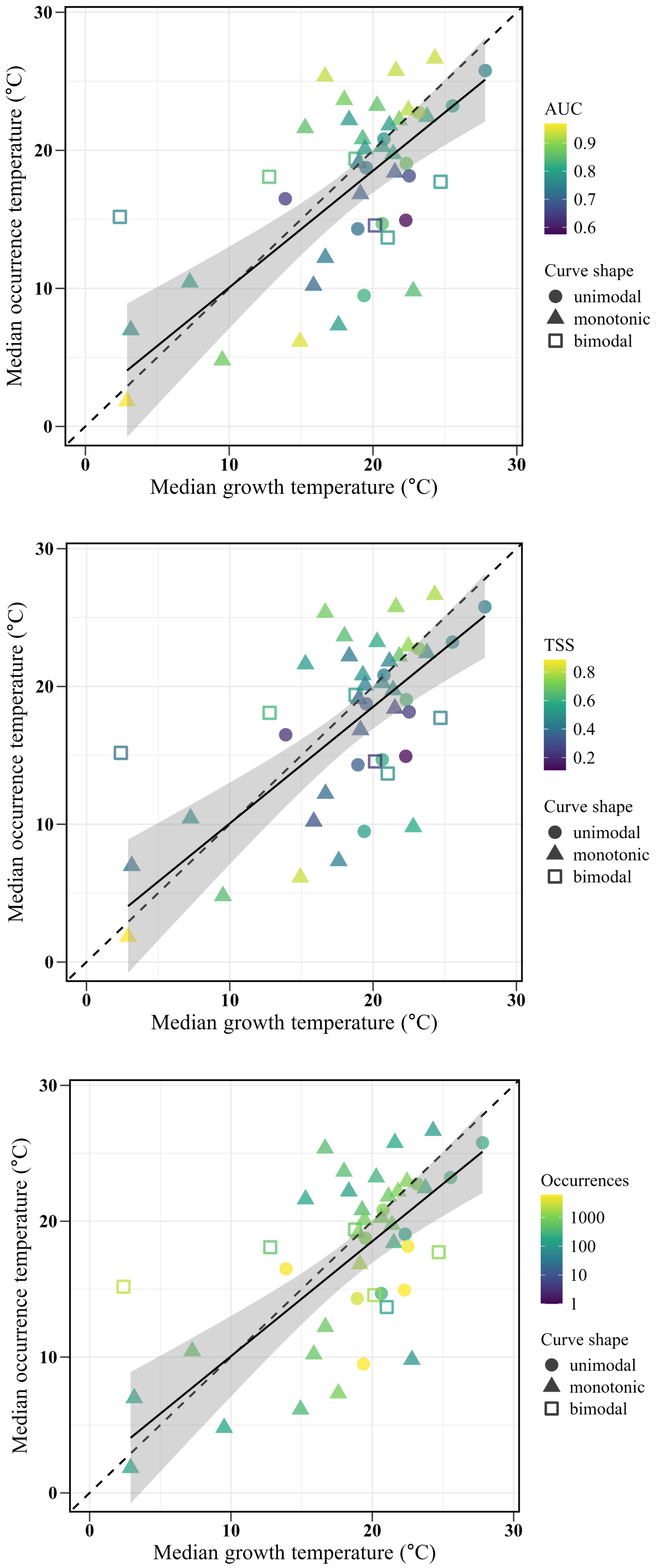

**Fig. S7**. The relationship between median growth and occurrence temperatures remains extremely similar if median growth temperature is calculated based on the mean of the individual strain estimates (slope = 0.86, intercept = 1.61, $R^2$ = 0.51, $p < 0.001$) instead of from the envelope, as we did in Fig. 2. The mean response (solid regression line) is indistinguishable from the 1:1 line (dashed line). The unimodal curves (circles) are likely most reliable, followed by monotonic (triangles) and finally bimodal (hollow squares) curves. The six bimodal curves were excluded from the regression, as we consider these estimates unreliable. Where lab growth curves exist for multiple strains of the same species, points represent the mean and error bars represent the range.

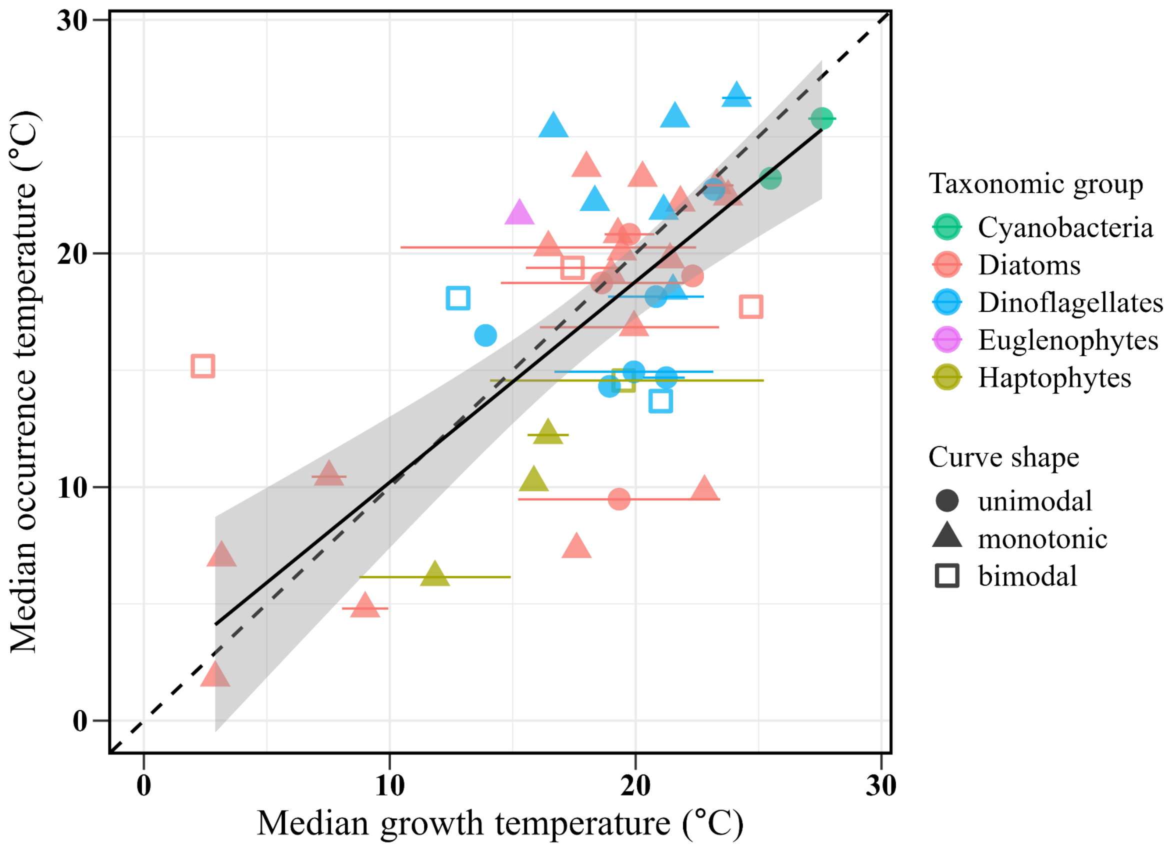

**Fig. S8**. The difference between median growth and median occurrence temperatures displays no clear relationship with growth niche width (slope = -0.06, intercept = 2.08, $p$ = 0.78, $R^2$ = 0.002). Positive values indicate higher median growth temperatures than median occurrence temperatures. Each point represents a species. The six species with bimodal occurrence curves (hollow squares) were excluded from the regression, as we consider their estimates unreliable. Where lab growth curves exist for multiple strains of the same species, species' median growth temperatures and growth niche widths were calculated from the envelope of strains' growth curves (Fig. S1).

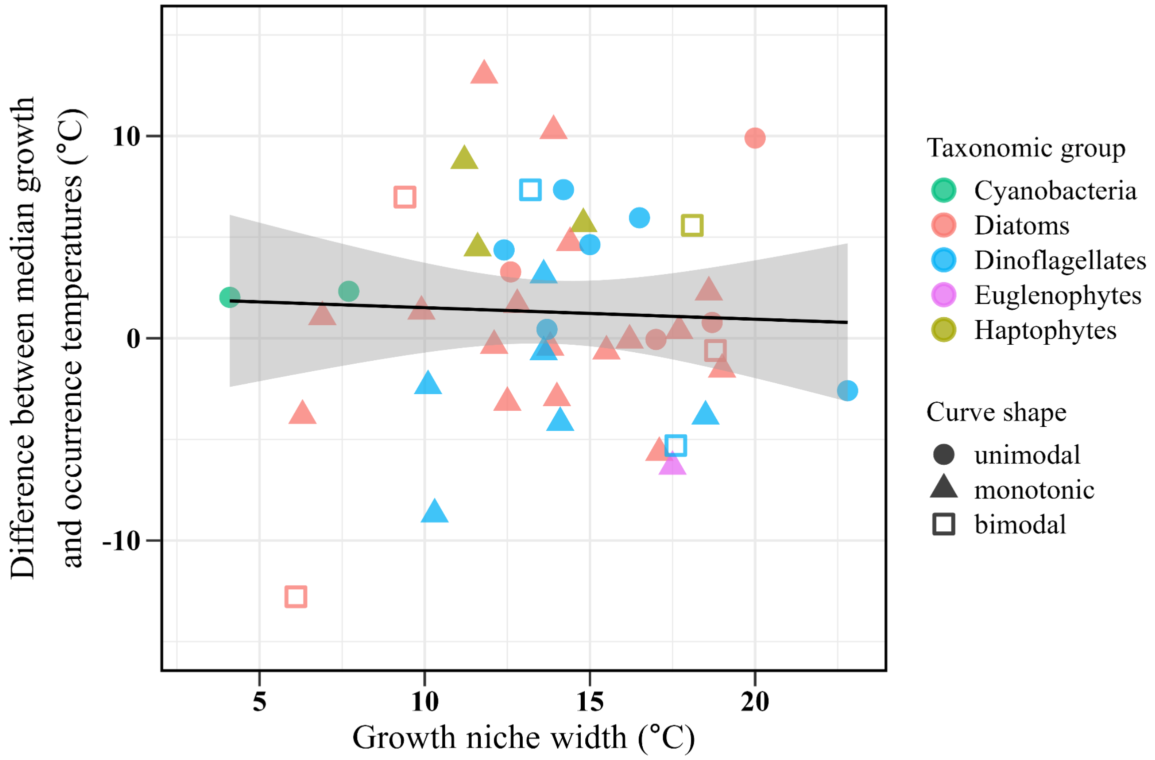

**Fig. S9**. The difference between species' median growth and occurrence temperatures displays no clear relationship with maximum growth rate across 39 marine phytoplankton species (slope = 0.81, intercept = 0.48, $p$ = 0.45, $R^2$ = 0.02). Positive values indicate higher median growth temperatures than median occurrence temperatures. Each point represents a species. The six species with bimodal occurrence curves (hollow squares) were excluded from the regression, as we consider their estimates unreliable. Where lab growth curves exist for multiple strains of the same species, species' median growth temperatures and maximum growth rates were calculated from the envelope of strains' growth curves.

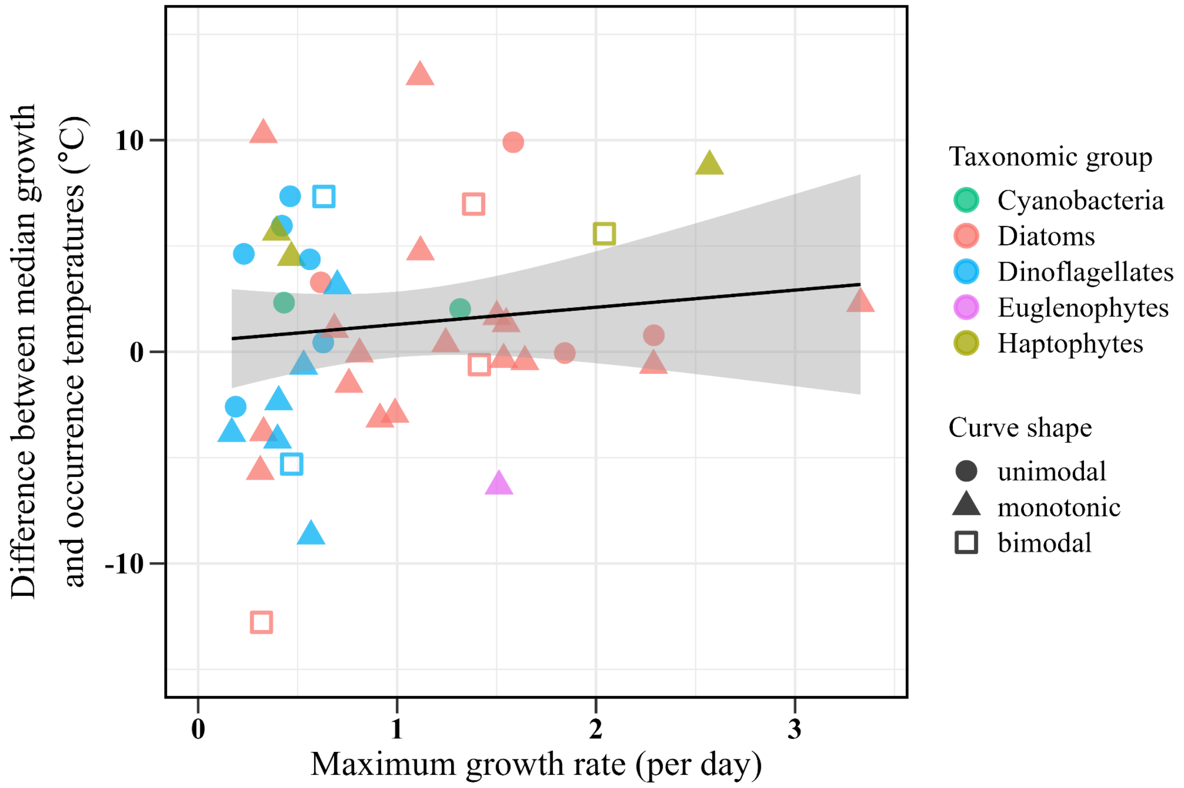

**Fig. S10**. The relationship between growth and occurrence niche widths is similar if growth niche width is calculated based on the mean of the individual strain estimates (slope = 0.55, intercept = 9.60, $p = 0.008$, $R^2 = 0.18$). The residual distribution exhibits mild deviation from normality but robust regression returns very similar coefficients and t-value estimates, so we present the OLS results here. The unimodal curves are likely most reliable, followed by monotonic and finally bimodal curves. The six bimodal curves (hollow squares) were excluded from the regression, as we consider these occurrence niche width estimates unreliable. Where lab growth curves exist for multiple strains of the same species, points represent the mean and error bars represent the range.

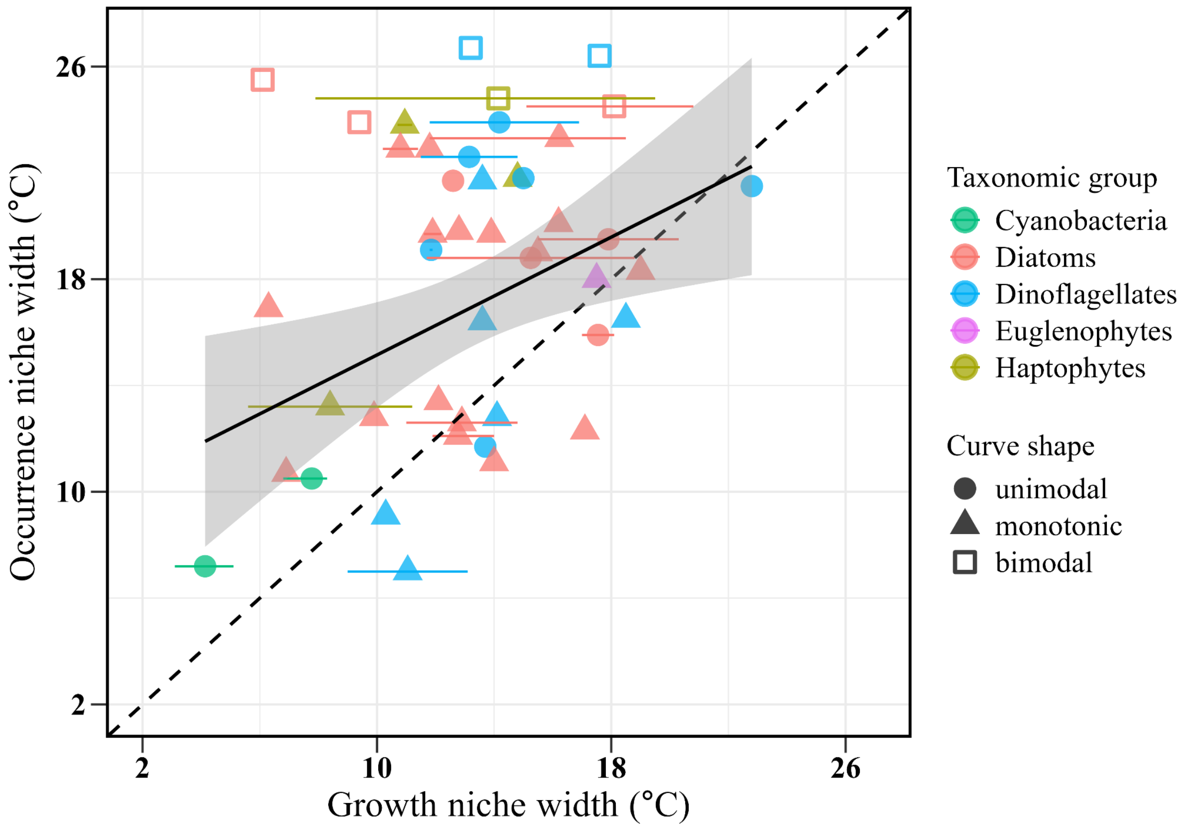
